# Morphometric Analysis of the Thymic Epithelial Cell (TEC) Network Using Integrated and Orthogonal Digital Pathology Approaches

**DOI:** 10.1101/2024.03.11.584509

**Authors:** Maria K. Lagou, Dimitrios G. Argyris, Stepan Vodopyanov, Leslie Gunther-Cummins, Alexandros Hardas, Theofilos Poutahidis, Christos Panorias, Sophia DesMarais, Conner Entenberg, Randall S. Carpenter, Hillary Guzik, Xheni Nishku, Joseph Churaman, Maria Maryanovich, Vera DesMarais, Frank P. Macaluso, George S. Karagiannis

## Abstract

The thymus, a central primary lymphoid organ of the immune system, plays a key role in T cell development. Surprisingly, the thymus is quite neglected with regards to standardized pathology approaches and practices for assessing structure and function. Most studies use multispectral flow cytometry to define the dynamic composition of the thymus at the cell population level, but they are limited by lack of contextual insight. This knowledge gap hinders our understanding of various thymic conditions and pathologies, particularly how they affect thymic architecture, and subsequently, immune competence. Here, we introduce a digital pathology pipeline to address these challenges. Our approach can be coupled to analytical algorithms and utilizes rationalized morphometric assessments of thymic tissue, ranging from tissue-wide down to microanatomical and ultrastructural levels. This pipeline enables the quantitative assessment of putative changes and adaptations of thymic structure to stimuli, offering valuable insights into the pathophysiology of thymic disorders. This versatile pipeline can be applied to a wide range of conditions that may directly or indirectly affect thymic structure, ranging from various cytotoxic stimuli inducing acute thymic involution to autoimmune diseases, such as myasthenia gravis. Here, we demonstrate applicability of the method in a mouse model of age-dependent thymic involution, both by confirming established knowledge, and by providing novel insights on intrathymic remodeling in the aged thymus. Our orthogonal pipeline, with its high versatility and depth of analysis, promises to be a valuable and practical toolset for both basic and translational immunology laboratories investigating thymic function and disease.

## INTRODUCTION

The thymus is a primary lymphoid organ, in which T cell development occurs^1^. The thymic environment subjects developing T cells, also known as thymocytes, to a series of crucial and intricate steps, necessary for their continued growth and competence. To support an assortment of specialized and unique developmental functions^2–4^, the thymus is compartmentalized into an outer cortical and an inner medullary zone. In general, the cortex oversees the homing, survival, and commitment of bone-marrow-derived early thymic progenitors (ETPs) to the T cell lineage. It also provides the requisite signals for the development of functional T cell receptors (TCR) through the process of positive selection, and promotes the proliferation/expansion of positively-selected thymocytes, promoting them into single-positive (SP) thymocytes expressing either CD4^+^ (SP4) or CD8^+^ (SP8)^2,4,5^. The medulla serves equally fundamental roles, such as the establishment of central tolerance and the development of regulatory T cells (T_regs_)^2,4,5^. The above processes are orchestrated via a complex, joint effort of an intrathymic stromal cell network, consisting of mesenchymal, endothelial, neuroendocrine, and dendritic cells, although the main protagonists are the highly specialized thymic epithelial cells (TECs)^4,6–9^.

Like other lymphoid organs, the thymus relies heavily on its distinct zonation to function properly^10,11^. Recent evidence has revealed that the traditional view of corticomedullary compartmentalization oversimplifies the complex nature of thymic environments, which includes additional, highly-specialized sub-niches. A prime example is a unique medullary sub-niche, known as the “perivascular space” (PVS), which was originally overlooked for its functional significance due to its appearance as an “epithelium-free area” in histological representations^12,13^. The entry of positively-selected single-positive (SP) thymocytes into the medulla triggers an interaction with AutoImmune Regulator^+^ (AIRE^+^) mTEC^hi^ (CD80^hi^, MHCII^hi^) and post-AIRE mimetic cells, crucial for establishing self-tolerant T cells^14,15^. After passing through this test, self-tolerant CD4^+^ and CD8^+^ T cells emigrate from the medullary parenchyma into the peripheral blood, a process known as “thymocyte egress”^16^. This process is tightly regulated by a complex network of thymic stromal cells, including endothelial cell, pericytes, and fibroblasts, which together orchestrate chemokine networks, shaping the microanatomy of the perivascular space^17–23^. As our understanding on thymic function deepens, it therefore becomes increasingly clear that this lymphoid organ is much more intricate and contextually organized than previously believed.

Given its pivotal role in adaptive immunity, the thymus has been predominantly explored via cell population analyses, utilizing flow cytometry^7,24,25^. This approach, however, has largely neglected the organ’s spatiotemporal dynamics, which are crucial for comprehensive, quantitative studies. In response, the Societies of Toxicological Pathology^1,26–28^ have provided the International Harmonization of Nomenclature and Diagnostic Criteria (INHAND) histopathology guidelines that, while valuable, are subjectively interpreted and lean towards a less objective framework. This gap in precise and universally accepted assessments hampers our understanding, especially as various pathologies can significantly alter thymic histological structure^27,28^. To address these issues, here, we have developed an orthogonal digital pathology pipeline that spans from microanatomical to microanatomical, and ultimately to ultrastructural analyses. Our innovative approach is anchored in a thoroughly vetted mouse model, chosen for its well-documented portrayal of age-related thymic involution. Through this proof-of-concept, we demonstrate the pipeline’s potential in meticulously charting thymic alterations across a spectrum of pathological conditions, offering a new lens through which to examine intrathymic remodeling.

## METHODS

### Animal Subjects

#### Ethics Statement

All studies involving mice were conducted in accordance with NIH regulations concerning the care and use of experimental animals with the approval of IACUC of Molecular Imaging, Inc. (Ann Harbor, MI), a facility accredited by the Association for Assessment and Accreditation of Laboratory Animal Care (AAALAC), or with the approval of Albert Einstein College of Medicine Animal Care and Use Committee.

#### Mice

Wild-type female FVB/NJ mice were acquired from NCI at 5-6 weeks of age, housed, and maintained in groups of 5 animals. After a post-arrival adaptation of 1-week, the mice were introduced to an “aging protocol”, during which they underwent weekly clinical and behavioral assessments, including body weight and pain assessment, based on an established grimace scale^29,30^. None of the mice fulfilled any endpoint criteria that could be regarded as confounding conditions in our study. Mice were humanely sacrificed as soon as they reached pre-specified age endpoints, and classified as “Young-Adult” (1.5-months, n=5), “Adult” (4-8 months, n=10), and “Old” (>18-months, n=11), into which they were allocated randomly. Euthanasia was performed after isoflurane anesthesia coupled to cervical dislocation. Mice housed under this aging protocol were treated identically until reaching the endpoint criteria.

#### Mouse Gender and Strain Selection

Due to established gender-related differences in the development of T cells^31–34^, it is generally not recommended to pool together male and female mice in studies related to thymic functions. When compared to males for example, aged female mice tend to have higher numbers of TEC, including the AIRE^+^ mTEC subpopulation, which is critical for the process of negative selection^31,33^. For purposes of method development, the current study has included female mice only. However, our proposed readouts can be performed in mice of any sex, age, strain, and body condition, without limitations. Finally, for evaluating strain-dependent alterations during aging, three 24-month-old C57Bl/6J mice, provided by the Maria-Cuervo lab, were included in the “Old” group, in addition to the FVB/NJ mice.

#### Necropsy Procedures

Complete necropsies were conducted on sacrificed mice, according to standardized protocols^35,36^, and age-associated lesions were recorded. Before sacrificing, whole body weight measurements were documented, and gross examinations were conducted on the extracted thymus and spleen after trimming peri-thymic and peri-splenic adipose tissue, respectively. The organs were weighted, fixed into 10% neutral buffered formalin (Fisher Scientific) for 24-48 hours, and paraffin embedded (FFPE). The weights were reported both as absolute and as “relative” values, normalized to body weights [Thymus Weight Index (TWI); Spleen Weight Index (SWI)].

### Histology and Immunofluorescence Staining, and Image Acquisition

#### Tissue Preparation

Unless otherwise stated, all slides, 5μm thick, were first dewaxed twice in xylene (Fisher Scientific) for 10min, then serially dehydrated twice in 100° ethanol (Fisher Scientific) for 10min, twice in 95° ethanol (Fisher Scientific) for 10min, and once in 70° ethanol (Fisher Scientific) for 10min, and finally rehydrated in distilled water for 5min.

#### Hematoxylin & Eosin (HE) Staining

All slides were stained using modified Meyer’s hematoxylin (Fisher Scientific) for 5min, then differentiated in 1% HCl, 70% ethanol (Fisher Scientific) for 5sec, then re-stained using Eosin Y (Fisher Scientific) for 1min 45sec, then cleared (95° ethanol, 100° ethanol, and xylene), and finally mounted with Permount (Fisher Scientific) medium.

#### Immunofluorescence Staining

Heat-Induced Antigen Retrieval (HIER) was performed with Citrate Buffer, pH 6.0 (Novus Biologicals) or EDTA, pH 8.0 (Epredia) in a steamer for 21min, followed by incubation with blocking solution (Goat serum, dilution 5:100 in Phosphate Buffered Saline Tween20,PBST, 0.05%, Fisher Scientific) for 1hour at room temperature. Primary antibody staining was performed overnight at 4°C, using: anti-Keratin-5 (Chicken anti-mouse; polyclonal; dilution 1:500 in blocking solution; Invitrogen); anti-Keratin-8 (Guinea Pig anti-mouse; polyclonal; dilution 1:100 in blocking solution; Acris); anti-AIRE (Rat anti-mouse; monoclonal: 5H12 clone; dilution 1:500 in blocking solution; Invitrogen); anti-DCLK1 (Rabbit anti-mouse; monoclonal: JA11-03 clone; dilution 1:200 in blocking solution; Invitrogen); anti-CD3 (Rabbit anti-mouse; monoclonal: SP7 clone; dilution 1:100 in blocking solution; Abcam); and anti-B220 (Rat anti-mouse; monoclonal: RA3-6B2 clone; dilution 1:100 in blocking solution; R&D systems). Secondary antibody staining was performed for 50min at room temperature, using: Goat anti-Rabbit Alexa488 (Invitrogen); Goat anti-Chicken Alexa488 (Invitrogen); Goat anti-Guinea Pig Alexa555 (Invitrogen); Goat anti-Rabbit Alexa555 (Invitrogen); Goat anti-Rat Alexa555 (Invitrogen); and Goat anti-Rat Alexa647 (Invitrogen), all diluted at 1:200 in blocking solution. Nuclear counterstaining was performed for 5min with 4′,6-diamidino-2-phenylindole (DAPI, dilution 1:1000 in PBST 0,05%; Novus Biologicals), or with Deep Red Anthraquinone 5 (DRAQ5, dilution 1:500 in PBST 0,05%; Invitrogen), before mounting with ProLong Gold anti-fade reagent (Invitrogen).

#### Microscopy and Slide Scanning

After completion of the HE or IF staining as described above, the slides were kept in the dark at room temperature, until imaged on a 3D HISTECH P250 Flash III automated slide scanner, using a 20x 0.75NA objective lens. The instrument automatically focuses and stitches the images into one single image file. For the IF in particular, each channel was acquired separately, with a set exposure time, and then combined into a multichannel overlay image. Further image analyses and development of morphometries were conducted in CaseViewer, ImageJ, and QuPath, as discussed below.

### Morphometry 1 – Corticomedullary Ratio

We developed three independent, quantitative, orthogonal methods, by using CaseViewer, ImageJ, and QuPath software^37^. Specifically, 5um thick, FFPE thymic tissue slides were stained with HE, or co-stained with DAPI/DRAQ5, Keratin-8 (KRT8) and Keratin-5 (KRT5), and then subjected to automatic slide scanning, as indicated above.

For <u>W</u>hole-<u>T</u>hymus methods, i.e., CMR_WT_, whole thymus IF images, depicting both lobes at their maximum width are captured using CaseViewer v2.4 (3D HISTECH). The images are exported as TIFF format, imported in QuPath v0.5.0, and the “image type” is set to “Fluorescence”. Three regions of interest (ROIs) are designated: (i) A “Whole-Thymus ROI”, corresponding to either single or double-positive KRT5/KRT8 signal; (ii) A “Medulla ROI” (i.e., medullary parenchyma) by demarcating the KRT5^+^ region only, and (iii) A “Cortex ROI” (i.e., cortical parenchyma), by masking the “Medulla ROI” out of the “Whole-Thymus ROI”, using an XOR logic gate. A similar approach is adapted for the HE stain. In this case, the “image type” is set to “Brightfield HE”. The cortex is distinguished from the medulla, based on its strong, basophilic stain, verified by two veterinary pathologists (A.H. and T.P.). The surface areas of the above ROIs are recorded in μm^>2^ and the CMR_WT_ is calculated for each image, using equation #1:

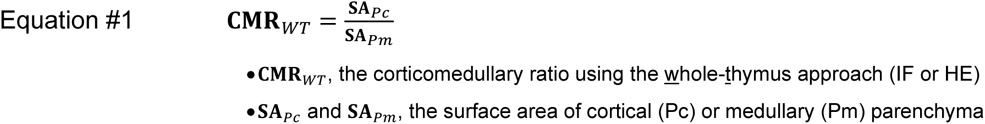

For the <u>T</u>hymic-<u>L</u>obule method, i.e., CMR_TL_, up to ten representative fields depicting single thymic lobules, are captured in HE stains, using CaseViewer v2.4. The fields are exported in TIFF format and processed in ImageJ v1.53q. One-dimensional ROIs of the cortical sheaths enveloping the intermediary medullary compartment in each lobule are then drawn in perpendicular fashion to the orientation of the corticomedullary border. These “straight line” ROIs range from one interlobular trabeculum to its opposite counterpart. The widths of the one-dimensional cortical and medullary ROIs are recorded in μm, and the CMR_TL_ is then calculated for each thymic lobule using equation #2:

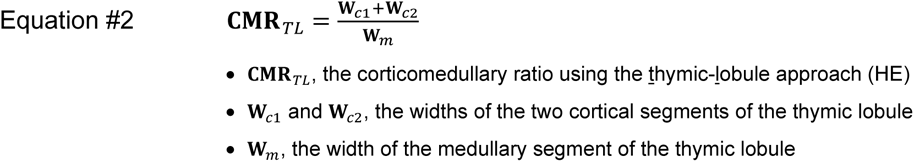

### Morphometry 2 – Topological Features of the cTEC Network

We developed multiple morphometric features to characterize the topology of the cTEC meshwork, using CaseViewer, ImageJ, and QuPath software^37^. Specifically, 5µm thick, FFPE thymic tissue slides were co-stained with DAPI, KRT8 and KRT5, and subjected to automatic slide scanning, as indicated above. Up to ten representative high-power-fields (HPFs) depicting the cortical parenchyma are captured using CaseViewer v2.4 (3D HISTECH), and exported as TIFF images for further analysis. To develop the cTEC network morphometries, the KRT8^+^ signal corresponding to the cTEC meshwork undergoes a sequence of image modifications: First, a “binary mask” is generated, based on intensity thresholding of the KRT8^+^ signal in 40x high power fields (HPF). Second, an “outline” of the binary mask corresponding to the borders of the KRT8^+^ signal is created. Third, using 63x derivatives of the previous images, KRT8^+^ objects are removed via the “Remove Outliers” tool in ImageJ, using a 2-pixel cutoff, while the remaining KRT8^+^ mask is subjected to the Skeletonize plugin^38^.

The first morphometry, <u>V</u>olume Index of cTEC (VI_cTEC_), reports the relative volume of the cTEC network as % fraction of the parenchymal volume of the cortex. Two specific ROIs are generated from the binary mask described above: (i) A “cTEC-ROI”, corresponding to KRT8^+^-only signal, and (ii) A “Cortex ROI”, corresponding to cortical parenchyma. The surface areas are measured in μm^2^, and the VI_cTEC_ is calculated for each field independently, using equation #3. The feature is then reported as a mean of all fields for each mouse.

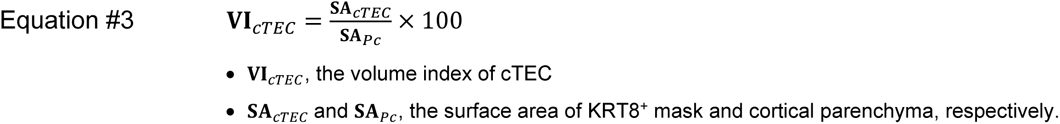

The second morphometry, Interface Index of cTEC (II_cTEC_), describes the surface area of the cTEC meshwork that is directly exposed for potential interaction to the thymocytes in the cortical parenchyma. The perimeter of the cTEC network is measured via the outline of the binary mask, described above, is expressed in μm, and is normalized to a given surface area unit of cortical parenchyma i.e., per 100μm^2^, using equation #4. The II_cTEC_ is calculated for each field independently and the feature is reported as a mean of all fields-of-view for each mouse.

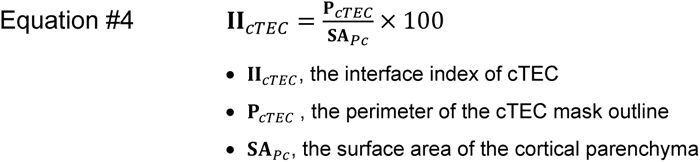

The third morphometry, <u>B</u>ranching Index of cTEC (BI_cTEC_), defines the branching features of the cTEC network, such as number of network branches, junctions, and average branch length. First, the skeletonized image generated above is analyzed without loop or end-point elimination using the Analyze Skeleton feature in ImageJ. The number of branches, junctions, and the average branch length, are collected as output from the analysis. The number of branches and junctions are normalized to a given surface area unit of cortical parenchyma, i.e., 100μm^2^, providing the BI1_cTEC_ and BI2_cTEC_ features, respectively, as shown in equations #5-6.

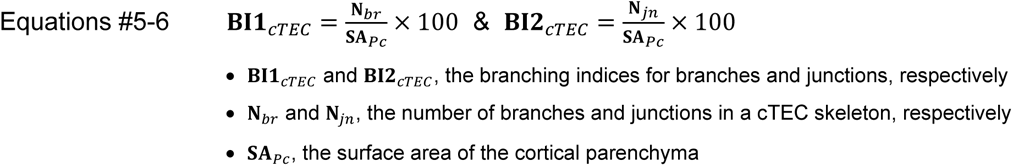

The average branch length, herewith abbreviated as BI3_cTEC_, is collected as a direct output from the skeleton analysis, and is expressed in μm per cTEC branch. It is calculated in each field-of-view, using equation #7.

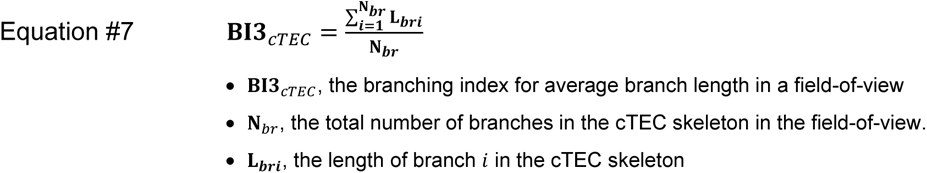

All three BIs are calculated for each field-of-view independently using equations #5-7, and the features are reported as means of all the fields-of-view for each mouse.

### Morphometry 3 – Perivascular Space (PVS)

The “Whole-Thymus” IF slides used for the calculation of corticomedullary ratios were also used to develop the PVS morphometry. Three regions of interest (ROIs) were thus designated: (i) A “Whole-Thymus ROI”, corresponding to either single or double-positive KRT5/KRT8 signal; (ii) A “Medulla ROI” (i.e., medullary parenchyma) by demarcating the KRT5^+^ region (which includes a prominent KRT8^+^ subset, as well), and (iii) The medullary regions that contain cells (DAPI^+^ or DRAQ5^+^), but do not harbor any mTEC (i.e., KRT5^+^KRT8^+^), characterize the epithelium-free PVS (i.e., KRT5^-^KRT8^-^), based on established knolwledge^17,39^. These PVS zones are demarcated and consolidated into a single “Merged-PVS ROI” for each field-of-view, using the OR logic gate. The surface areas of all the above ROIs are recorded in μm^2^ for each field-of-view, independently.

Two morphometric features are reported. The first feature reports the fraction of PVS (%) occupying the medullary parenchyma (PVS_Pm_), while the second feature reports the fraction of the PVS corresponding to the surface area of the whole thymus (PVS_WT_). Both features are first calculated in a single field-of-view, using equations #8-9), and then reported as means of all fields-of-view for each mouse.

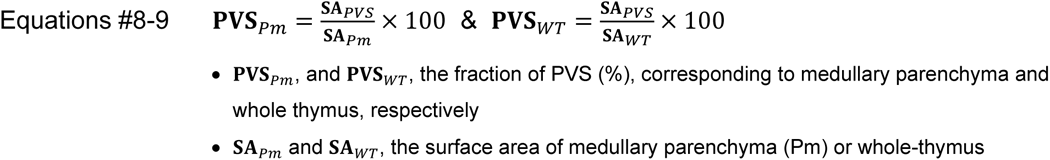

### Morphometry 4 – Topological Features of the mTEC Network

The IF-stained thymic tissue slides used above were also used for mTEC analysis. Up to ten representative 20x magnification images of thymic medulla were captured, then exported as TIFF images, and analyzed on ImageJ v1.53q. The following regions of interest (ROIs) were then designated: (i) A “Medulla-ROI” (i.e., medullary parenchyma) by demarcating the KRT5^+^ region only, (ii) The “Merged-PVS ROI”, as indicated above, and (iii) The “Epithelium-Containing Area (ECA) ROI”, by masking the “Merged-PVS ROI” out of the “Medulla-ROI”.

Two morphometries were developed to characterize the mTEC network, the VI_mTEC_ and II_mTEC_, in similar analogy to the cTEC counterparts. In this case, the KRT5^+^ signal corresponding to the mTEC network undergoes the sequence of the first two modifications, thus omitting the skeleton analysis. First, a “binary mask” is generated, based on intensity thresholding of KRT5^+^ signal in 20x high power fields (HPF). Second, an “outline” of the binary mask corresponding to the borders of the KRT5^+^ signal is created. The two features are calculated in an analogous manner with only one difference. Instead of normalizing to the total surface of medullary parenchyma as in the case of the cTEC morphometries, the normalization occurs with the surface of the epithelium-containing area (i.e., “ECA-ROI”), because each medullary field-of-view has a diverse composition of PVS zones, and these may change with aging. As before, the two features are calculated in each field-of-view independently, using equations #10-11, and the features are reported as means of all fields-of-view for each mouse.

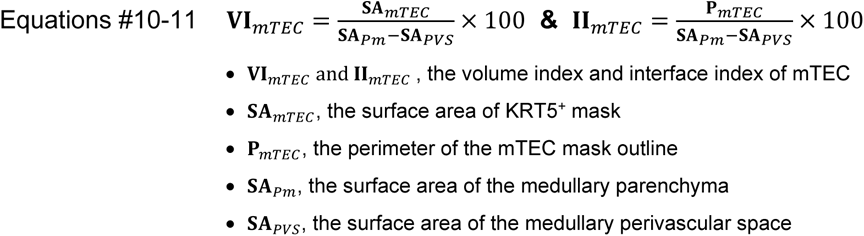

### Morphometry 5 – Topological Features of mTEC Subset Distribution

We developed morphometric features to characterize the spatial distribution of specific mTEC subsets, in particular the AIRE^+^ mTEC^hi^ and the Doublecortin-like kinase 1^+^ (DCLK1^+^) thymic tuft cells. Specifically, 5µm thick, FFPE thymic tissue slides were co-stained with DAPI, KRT5, AIRE, and DCLK1, and subjected to automatic slide scanning, as indicated above. Up to five representative high-power-fields (HPFs) depicting the medullary parenchyma were captured using CaseViewer v2.4 (3D HISTECH), and uploaded on QuPath v0.5.0^37^. The “Medulla ROI” was automatically demarcated, using an in-built pixel classifier with the following settings: moderate resolution; Gaussian prefilter; smoothing sigma 2.5; and threshold 7. This classifier successfully removes non-medullary areas displayed in the field-of-view, such as epithelium-free PVS zones, and KRT5^-^ thymic cortical segments. The AIRE^+^ and DCLK1^+^ cell populations were also automatically demarcated and counted, using the cell detection tool with the following settings for DCLK1^+^ cells: requested pixel size 0.5 μm; background radius 8μm, sigma 0.5μm; Min/Max Area 20-500μm^2^; and threshold 65. The respective settings for AIRE^+^ mTEC^hi^ subset were: requested pixel size 0.5 μm; background radius 8μm, sigma 3μm; Min/Max Area 10-150μm^2^; and threshold 35. Slight modifications to these settings may be made across different batches of staining, depending upon the quality/pattern of staining. Finally, areas corresponding to erythrocytes, dead cells, debris, cysts, and other artifacts, are manually removed, using the brush/eraser tool.

The first morphometric feature is the “density” (d) of the AIRE^+^/DCLK1^+^ mTEC cell subsets. This morphometry reports the number of AIRE^+^/DCLK1^+^ mTEC cells in a given surface area unit (mm^2^) of epithelium-containing medullary parenchyma. The feature is calculated in one field-of-view, using equations #12-13, and is reported as a mean of all fields-of-view for each mouse.

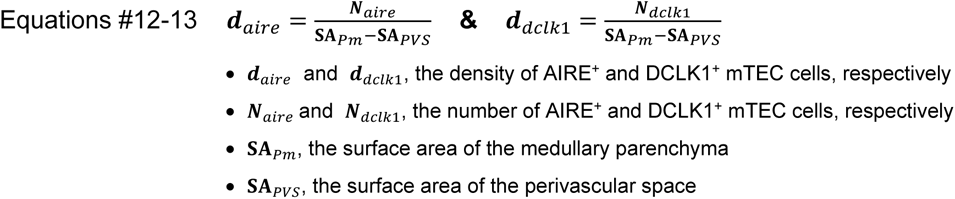

The second morphometric feature of spatial distribution is the “homogeneity” (h) of spatial arrangement of the AIRE^+^/DCLK1^+^ cell subsets. To calculate this feature, a series of modifications on the already obtained fields-of-view is performed, immediately following identification of the “Medullary ROI” using the pixel classifier, and counting of the cells of interest using the cell detection tool, as described above. First, a “template ROI” is applied to the image, to segment the “medullary ROI” into 20 equally-sized, rectangular-shaped sectors. Because up to 5 fields-of-view per animal are retrieved, the total number of sectors may thus vary between 20 and 100 per animal. The density of the AIRE^+^/DCLK1^+^ mTEC cells is then independently calculated for each sector *i*, using equations #14-15.

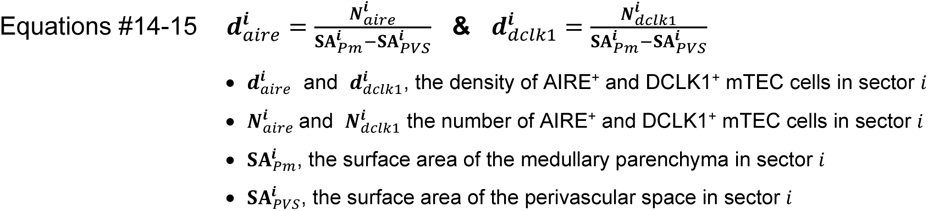

The standard deviation of sector densities of AIRE^+^/DCLK1^+^ mTEC cells is then calculated for a reference group (SD^ref^), across all fields-of-view, using equations #16-17. We selected the Young-Adult as the reference group, because this group has the most structurally and functionally active thymus, after which involution begins, and as such, we assumed that any deviations observed within the Young-Adult thymus should possess physiological/biological relevance.

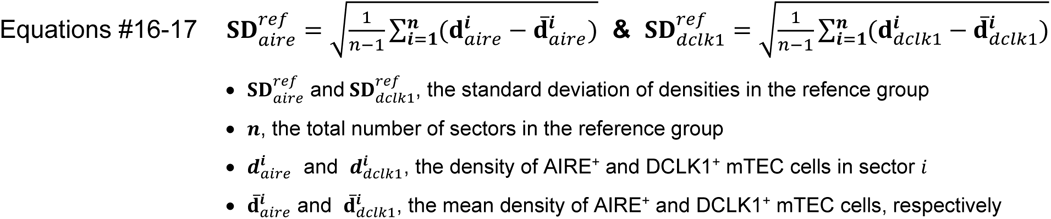

Using data from equations #14-17, each sector is subjected to the following logical checks, independently, and the number N of TRUE returns is counted for each mouse:

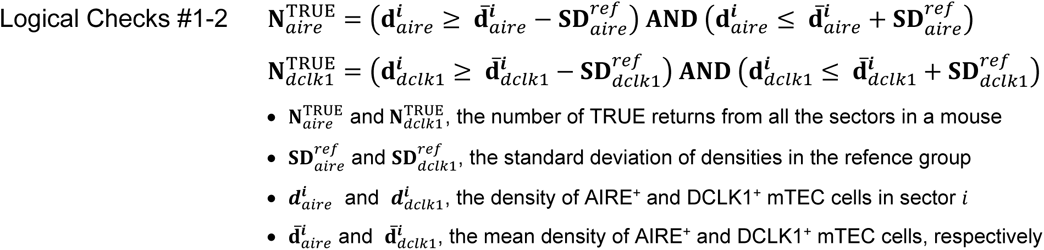

These logical checks #1-2 will return TRUE if the sector’s density falls within 1 SD^ref^ from the mean density, and FALSE otherwise. Following these determinations, the homogeneity index, i.e., **h***_aire_* and **h***_dclk1_*, reports the fraction of sectors that checked TRUE on these logical checks, for each mTEC subset of interest, using equations #18-19.

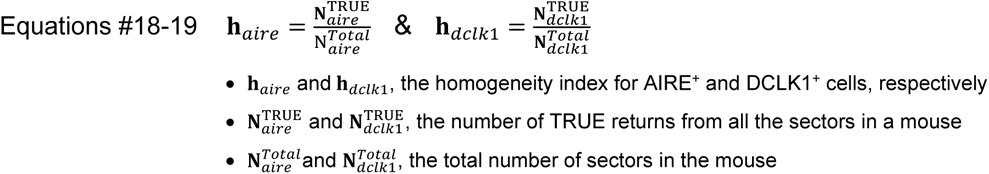

The homogeneity index shows the fraction of sectors, whose density distribution falls within an expected variance (i.e., standard deviation of a reference group). A high **h** index informs that most sectors (i.e., medullary micro-niches) tend to have similar densities of specific mTEC cell subsets. A low **h** index, however, informs that a large number of thymic micro-niches presents with either very-high or very-low density of specific mTEC cell subsets, thus implying that the overall spatial distribution/arrangement within the medulla is more heterogeneous and clustered.

### Morphometry 6 – White-to-Red Pulp Ratio in the Spleen

We developed two independent methods for calculating White-to-Red Pulp Ratio (WRPR) in the spleen, using CaseViewer and QuPath. Spleen tissue slides were stained with either HE, or multiplex IF consisting of DAPI, CD3 (to label the T cells), and B220 (to label the B cells), and subjected to automatic slide scanning, as described above. Whole spleen images depicting the splenic parenchyma surrounded by an intact splenic capsule, are captured using CaseViewer v2.4, and exported as TIFF images for further analysis in QuPath v0.5.0.

For the IF-stained slide, the “image type” is set to “Fluorescence”. Three regions of interest (ROIs) are designated: (i) A “Whole-Spleen ROI”, which is demarcated by the splenic capsule to include the splenic parenchyma; (ii) A “White-Pulp ROI”, corresponding to strong CD3/B220 signal exhibiting follicular organization; (iii) A “Red-Pulp ROI”, by masking the “White-Pulp ROI” out of the “Whole-Spleen ROI”, using the XOR logic gate. A similar approach was adapted for calculating the ratio, using an HE stain. In this case, the “image type” is set to “Brightfield HE” in QuPath, and the white pulp is distinguished from the red pulp, exclusively based on the strong, basophilic stain of the lymphoid follicles, and is confirmed by veterinary pathologists (A.H. and T.P.). The surface areas of the above ROIs are recorded in μm^2^ and the WRPR_WS_ is calculated for each image, using equation #20.

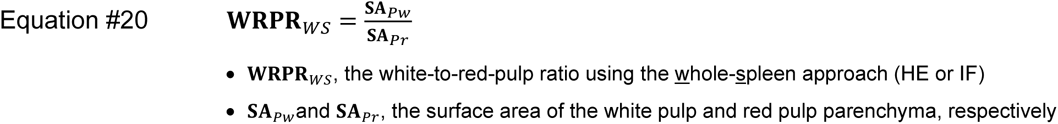

### Electron Microscopy

#### Tissue Preparation

Fresh thymic tissues were processed for Transmission Electron Microscopy (TEM) (n=7), or with an Osmium-Thiocarbohydrazide-Osmium (OTO) method (n=2), for 3D-SEM Array Tomography, as will be discussed below.

#### Transmission Electron Microscopy (TEM)

Samples were fixed with 2.5% glutaraldehyde, 2% paraformaldehyde in 0.1 M sodium cacodylate buffer, postfixed with 1% osmium tetroxide followed by 2% uranyl acetate, dehydrated through a graded series of ethanol and embedded in LX112 resin (LADD Research Industries, Burlington VT). Ultrathin sections were cut on a Leica Ultracut UC7, stained with uranyl acetate followed by lead citrate and viewed on a JEOL 1400 Plus transmission electron microscope at 120kV.

#### OTO Processing and Scanning Electron Microscopy (OTO-SEM)

Samples were immersion fixed in 2.0% paraformaldehyde, 2.5% glutaraldehyde in 0.1M sodium cacodylate buffer, then processed using a modified National Center for Microscopy and Imaging method of OTO^40^. In brief, samples were post fixed with reduced osmium, treated with thiocarbohydrazide, further stained with osmium, *en bloc* stained with uranyl acetate, further stained with lead aspartate, dehydrated in a graded series of ethanol, and embedded into LX112 resin. 55nm-thick sections were cut on a Leica Artos microtome using a Diatome AT 35° knife, and picked up on freshly glowed silicon wafers. Sections were observed on Zeiss Supra 40 Field Emission Scanning Electron Microscope in backscatter mode, using an acceleration voltage of 8.0 kV.

#### Three-Dimensional Reconstruction of the cTEC Meshwork

Regions of interest were collected with ATLAS 5.0, using a pixel size of 6.0 x 6.0 nm and a dwell time of 6.0 µs. Images were aligned and segmentation was done using the IMOD suite of programs, as described^41,42^. Briefly, stacks of images were aligned in IMOD using Midas and cells of interest were manually segmented out using the drawing tools in 3DMOD. cTEC cells were identified by their large, euchromatic nucleus, nucleolus shape, and the presence of tonofilaments throughout the cell.

### Statistical Analysis

#### Statistical Package

GraphPad Prism 10 software was used for generation of all graphs and for statistical hypothesis testing.

#### Statistical Hypothesis Testing

Results are presented as hybrid scatterplot/bar graphs, with means and Standard Error of the Mean (SEM). Linear correlations are presented as correlation matrices or scatterplots with simple linear regression lines. Comparison of variables among age groups is performed with One-Way ANOVA and post-hoc analysis using the Dunnett’s multiple comparisons test. Linear correlations are analyzed using Spearman’s correlation and coefficient of determination (r). For all graphs, *p* values of ≤0.05 are stated as statistically significant, while *p* values >0.05 are as non-significant (ns). For all graphs, (*) corresponds to *p*≤0.05, (**) corresponds to *p*<0.01, (***) corresponds to *p*<0.001, and (****) corresponds to *p*<0.0001.

#### Exclusion Criteria for Individual Cases

All datasets underwent an outlier calculator testing using an alpha of 0.05. Any values falling outside the 95% confidence interval in specific variables were excluded. Individual cases were also excluded if they lacked sufficient thymic or splenic tissue, to generate enough or representative regions of interest (ROIs) for analysis.

#### Blindness in Image Analysis

All pathologists involved in histopathological diagnoses and quantifications were blinded to the age group allocations. Scientists performing AIRE^+^ mTEC homogeneity analysis, AIRE^+^ and DCLK1^+^ mTEC density analyses, and electron microscopy 3D reconstructions, were also completely blinded. All other variables were calculated by scientists that were not blinded to age group allocation. However, to eliminate all the operator-dependent biases, all the samples were treated and processed in the same way, i.e., snapshots from each case were captured and analyzed together in one batch, using the same settings (e.g., intensity thresholding, etc.) in the associated digital pathology software.

## RESULTS

### Mouse Cohort Necropsy – General Findings and Progressive Age-Associated Lesions

To develop a digital pathology framework for assessing thymic lesions, we employed a mouse model of age-dependent thymic involution, in which mice were sacrificed at diverse age groups, and experimental endpoints were assessed **(Fig. 1A)**. As expected, we found that the total body weight of the mice significantly increased over time **(Fig. 1B)**. Because it is typical for aged mice to confer spontaneous age-associated lesions, we performed whole body necropsy, to determine the possibility of unreported pathologies that could bias our study. In general, we found no severe pathologies in the Young-Adult and Adult mouse groups. However, certain Old mice developed spontaneous lesions, diagnosed by veterinary pathologists **(Fig. S1A-F; Table S1)**: Multifocal mild-to-moderate neutrophilic, histiocytic, lymphoplasmacytic, necrotising steatitis with mineralisation and hemosiderin-laden macrophages in the omentum **(Figs. S1A-A’)**; Adrenal gland adenoma type B and pheochromocytoma **(Figs. S1B-B’)**; Multifocal solid, bronchiolo-alveolar lung carcinoma **(Figs S1C-C’)**; Multifocal lymphocytic proliferation of the renal pelvis and associated vasculitis of the perirenal fat vessels **(Figs. S1D-D’)**; Bone marrow hyperplasia **(Figs. S1E-E’)**; Splenic hyperplasia **(Figs. S1F-F’)**. These lesions are typical and expected to spontaneously rise in mice during aging^43^. Because there is no literature linking any of these lesions to specific thymic defects, except that they may represent general age-associated stressors, we reasoned that our animal cohort was suitable for further analysis.

**Figure 1.**
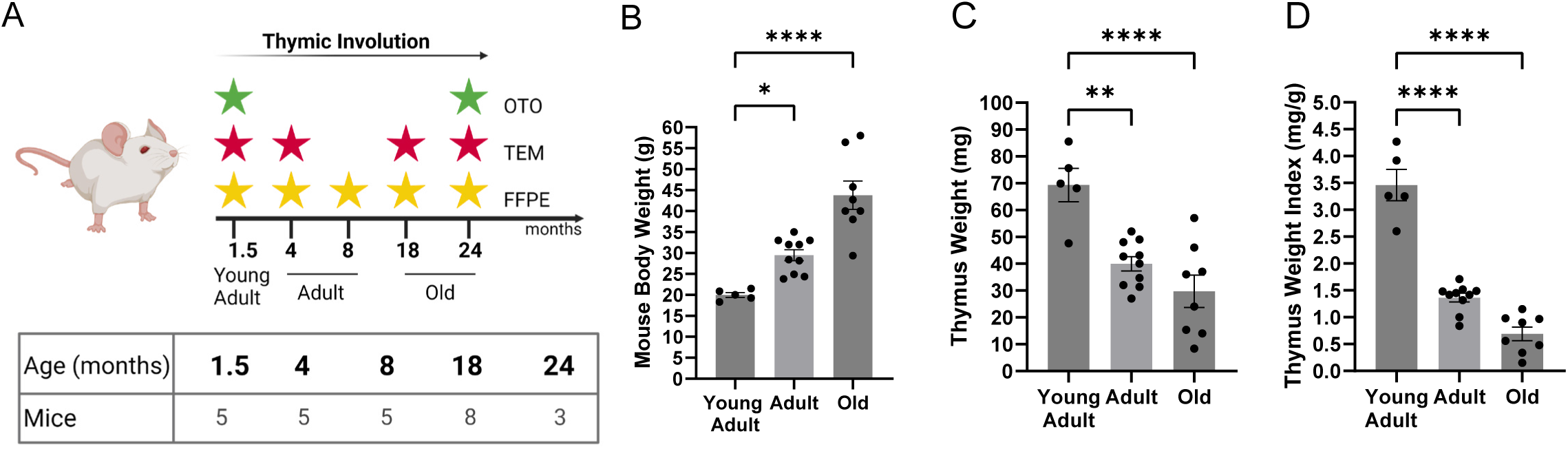
Mouse model of age-dependent thymic involution. **(A)** Experimental design and timepoints for assessing morphometric readouts. *Yellow star, Formalin-fixed, paraffin embedded (FFPE) hematoxylin & eosin (HE) and immunofluorescence (IF) analysis; red star, transmission electron microscopy (TEM); green star; osmium-thiocarbohydrazide-osmium (OTO) coupled to 3D scanning electron microscopy (SEM) array tomography.* (**B)** Mouse body weight for each group shown in (A). **(C)** Absolute weight of the extracted thymi for each group shown in (A). **(D)** Thymus weight, normalized to body weight of each mouse shown in (A), i.e., thymus weight index (TWI). *One-Way ANOVA; *p*≤*0.05; **p*≤*0.01; ***p*≤*0.001; ****p*≤*0.0001*.

**Table 1:** Histopathological descriptions and diagnoses of spontaneous, incidental findings in the Old mouse group. The first column refers to the code assigned to each tissue with lesions.

| Mice - Lesions | Diagnosis | Microscopic description |
| --- | --- | --- |
| GSK_30_Abdominal masses_20 | Omentum; multifocal mild lymphoplasmacytic and histiocytic steatitis with multifocal extensive (coagulative) necrosis with multifocal mineralisation and haemosiderin laden macrophages | Omentum: Five sections composed of the omentum are examined. In the majority of the omental sections there are multifocal to coalescing small to moderate numbers of macrophages, hemosiderin laden macrophages, neutrophils and fewer lymphocytes and plasma cells surrounding and replacing adipocytes (crown-like structures). There are occasional areas of mineralisation and necrosis. There is no evidence of infectious agents or neoplasia in the examined sections. |
| GSK_30_Abdominal masses_21 | Adrenal gland, omentum and liver; favour malignant pheochromocytoma | Adrenal gland: Multiple sections are examined. In the majority of the sections, effacing and replacing the previous structure (presumed origin from medulla of the adrenal gland) is an unencapsulated, multinodular, infiltrative, densely cellular, malignant round cell neoplasm. Neoplastic cells are arranged in dense solid areas and packets frequently palisading around necrotic areas and supported by a prominent collagenous stroma. Individual cells are round with variably distinct borders, a clear cytoplasm, and round nuclei with coarsely stippled chromatin and 2-4 prominent nucleoli. Anisocytosis and anisokaryosis are moderate. There are 9 mitotic figures observed in 10 HPF 2.37 mm<sup>2</sup>. The neoplasm is multifocally replaced by large areas of coagulative necrosis. |
| GSK_30_Abdominal masses_22 | Omentum; multifocal mild to moderate neutrophilic, histiocytic, lymphoplasmacytic, necrotising steatitis with mineralisation and haemosiderin laden macrophages | Omentum: Eight sections, seven composed of the omentum and one entirely of muscle are examined. In the majority of the omental sections there are multifocal to coalescing small to moderate numbers of macrophages, hemosiderin laden macrophages, neutrophils and fewer lymphocytes and plasma cells surrounding and replacing adipocytes (crown-like structures). There are occasional areas of mineralisation and necrosis. There is no evidence of infectious agents or neoplasia in the examined sections. |
| GSK_30_ADR_21 | Adrenal gland; adenoma (type B) | Adrenal gland: One section composed of adrenal gland and omentum is examined. Expanding and replacing the adjacent cortex is a single well demarcated, densely cellular, benign epithelial neoplasm. Neoplastic cells are arranged in broad trabeculae and nests supported by a thin fibrovascular stroma. Individual cells are polygonal with distinct borders, a large amount of finely vacuolated eosinophilic cytoplasm and a large round nucleus with finely stippled basophilic chromatin and one prominent nucleolus. Anisocytosis and anisokaryosis are moderate and no mitotic figures are observed in 2.37mm<sup>2</sup>. The neoplasm is completely contained within the excised adrenal gland |
| GSK_30_ADR_22 | Adrenal gland; adenoma (type B) | Adrenal gland: Two sections composed of adrenal gland neoplasm are examined. Expanding and replacing the adjacent cortex are two well demarcated, densely cellular, benign epithelial neoplasms. Neoplastic cells are arranged in broad trabeculae and nests supported by a thin fibrovascular stroma. Individual cells are polygonal with distinct borders, a large amount of finely vacuolated eosinophilic cytoplasm and a large round nucleus with finely stippled basophilic chromatin and one prominent nucleolus. Anisocytosis and anisokaryosis are moderate and no mitotic figures are observed in 2.37mm<sup>2</sup>. |
| GSK_30_ADR_23 | Adrenal gland; adenoma (type B) | Adrenal gland: Seven sections composed of adrenal gland neoplasm, muscle and omentum are examined. Expanding and replacing the adjacent cortex are two well demarcated, densely cellular, benign epithelial neoplasms. Neoplastic cells are arranged in broad trabeculae and nests supported by a thin fibrovascular stroma. Individual cells are polygonal with distinct borders, a large amount of finely vacuolated eosinophilic cytoplasm and a large round nucleus with finely stippled basophilic chromatin and one prominent nucleolus. Anisocytosis and anisokaryosis are moderate and no mitotic figures are observed in 2.37mm<sup>2</sup>. |
| GSK_30_BM_21 | Bone marrow; hyperplasia | Bone marrow: Multiple bone and bone marrow samples are examined. In the bone marrow the cellularity is increased and granulocytic. Increased numbers of densely packed myeloid cells fill the medullary space, increasing the myeloid to erythroid (M:E) ratio. Megakaryocytes are present. Increased band cells identify granulocytic precursor cells. Response to a peripheral abscess and demand for neutrophils. (Differentiation of granulocytic hematopoietic hypercellularity from well-differentiated granulocytic leukemia is challenging.) |
| GSK_30_BM_22 | Bone marrow; hyperplasia | Bone marrow: Multiple bone and bone marrow samples are examined. In the bone marrow the cellularity is increased and granulocytic. Increased numbers of densely packed myeloid cells fill the medullary space, increasing the myeloid to erythroid (M:E) ratio. Megakaryocytes are present. Increased band cells identify granulocytic precursor cells. Response to a peripheral abscess and demand for neutrophils.(Differentiation of granulocytic hematopoietic hypercellularity from well-differentiated granulocytic leukemia is challenging.) |
| GSK_30_BM_23 | Bone marrow; hyperplasia | Bone marrow: Multiple bone and bone marrow samples are examined. In the bone marrow the cellularity is increased and granulocytic. Increased numbers of densely packed myeloid cells fill the medullary space, increasing the myeloid to erythroid (M:E) ratio. Megakaryocytes are present. Increased band cells identify granulocytic precursor cells. Response to a peripheral abscess and demand for neutrophils.(Differentiation of granulocytic hematopoietic hypercellularity from well-differentiated granulocytic leukemia is challenging.) |
| GSK_30_KIDN_20 | Kidney, renal pelvis; multifocal lymphocytic proliferation | Kidney: Two sections are examined. There is diffuse congestion in both sections. There are multifocal aggregates of monomorphic lymphocytes in the renal pelvis and within the cortex. There is no evidence of infectious agents or neoplasia in the examined sections. |
| GSK_30_KIDN_21 | 1. Perirenal fat vessel; multifocal lymphocytic proliferation and associated vasculitis<br>2. Kidney; multifocal occasional protein casts and mineral | Kidney and perirenal fat vessel: Two sections are examined. There is diffuse congestion in both sections. Within the kidney parenchyma in one section there are intratubular casts with protein and mineral and occasional dark brown pigment. There is diffuse congestion in both sections. surrounding a vessel in the perirenal fat there are multifocal aggregates of monomorphic lymphocytes and the associated vessel wall is expanded by a mixed infiltrate of lymphocytes, neutrophils admixed with cellular debris (favour vasculitis) . Within the perirenal fat there are multifocal crown like structures composed of macrophages , hemosiderin laden macrophages and fewer lymphocytes. There is no evidence of infectious agents or neoplasia in the examined sections. |
| GSK_30_LIV_20 | Liver; multifocal, moderate, hepatocellular, glycogen-type vacuolar degeneration | Liver: Two sections are examined. The hepatic lobular architecture is maintained, with an expected distance between portal triads. Within portal triads are an expected number of biliary profiles and the hepatic arteries are easily identifiable. Multifocally, randomly, hepatocytes are mildly to moderately distended with a small amount of intracytoplasmic, feathery vacuolation (glycogen-type vacuolar degeneration). There is no evidence of neoplasia or infectious agents in the sections examined. |
| GSK_30_LIV_22 | Liver; diffuse mild, hepatocellular, glycogen-type vacuolar degeneration | Liver: Two sections are examined. The hepatic lobular architecture is maintained, with an expected distance between portal triads. Within portal triads are an expected number of biliary profiles and the hepatic arteries are easily identifiable. Diffusely, hepatocytes are mildly distended with a small amount of intracytoplasmic, feathery vacuolation (glycogen-type vacuolar degeneration). There is no evidence of neoplasia or infectious agents in the sections examined. |
| GSK_30_LIV_23 | Liver; diffuse mild, hepatocellular, glycogen-type vacuolar degeneration with rare single cell necrosis | Liver: Two sections are examined. The hepatic lobular architecture is maintained, with an expected distance between portal triads. Within portal triads are an expected number of biliary profiles and the hepatic arteries are easily identifiable. Diffusely, hepatocytes are mildly distended with a small amount of intracytoplasmic, feathery vacuolation (glycogen-type vacuolar degeneration). There are rare areas of single cell necrosis with associated lymphoplasmacytic aggregates admixed with neutrophils. There is no evidence of neoplasia or infectious agents in the sections examined. |
| GSK_30_LUNG_20 | Lung; solid bronchiolo-alveolar carcinoma (3mm diameter) | Lung: Focally effacing the lung parenchyma is a well-demarcated, densely cellular, epithelial neoplasm. Neoplastic cells are arranged in single and multiple layers arising from the alveolar lining and/or small airway epithelium. Within the centre of the neoplasm the neoplastic cells occlude the alveolar spaces and form solid nests. Neoplastic cells have sharp cell borders, a moderate amount of brightly eosinophilic cytoplasm and one single, frequently pleomorphic nucleus with coarsely stippled chromatin. Anisokaryosis and anisocytosis are frequent. There are 14 mitoses in ten high power fields.<br>Thyroid gland: No significant histopathological findings.<br>Lymph node, mediastinal: Reactive mediastinal lymph node. |
| GSK_30_LUNG_22 | Lung; multifocal solid, bronchiolo-alveolar carcinoma (3.6mm diameter) | Lung: Multifocally effacing the lung parenchyma are two well-demarcated, densely cellular, epithelial neoplasms. Neoplastic cells are arranged in single and multiple layers arising from the alveolar lining and/or small airway epithelium. Within the centre of the neoplasm the neoplastic cells occlude the alveolar spaces and form solid nests. Neoplastic cells have sharp cell borders, a moderate amount of brightly eosinophilic cytoplasm and one single, frequently pleomorphic nucleus with coarsely stippled chromatin. Anisokaryosis and anisocytosis are frequent. There are 17 mitoses in ten high power fields.<br>There are multifocal aggregates of lymphocytes within the mediastinal fat and the lung parenchyma. There is multifocal eosinophilic crystalline pneumonia. |
| GSK_30_SPL_21 | Spleen, red pulp; extramedullary hematopoiesis, increased megakaryocytes with haemosiderosis | Spleen: One section is examined. Throughout the splenic parenchyma are numerous megakaryocytes and moderate numbers of erythroid and myeloid precursors with occasional mitotic activity (extramedullary haematopoiesis). The splenic architecture is maintained, with evident lymphoid follicles. There are multifocal increased numbers of hemosiderin macrophages. |
| GSK_30_SPL_22 | Spleen, red pulp; extramedullary hematopoiesis, increased megakaryocytes with haemosiderosis | Spleen: One section is examined. Throughout the splenic parenchyma are numerous megakaryocytes and moderate numbers of erythroid and myeloid precursors with occasional mitotic activity (extramedullary haematopoiesis). The splenic architecture is maintained, with evident lymphoid follicles. There are multifocal increased numbers of hemosiderin macrophages. |
| GSK_30_SPL_23 | Spleen, red pulp; extramedullary hematopoiesis, increased megakaryocytes with haemosiderosis | Spleen: Three section are examined. Throughout the splenic parenchyma are numerous megakaryocytes and moderate numbers of erythroid and myeloid precursors with occasional mitotic activity (extramedullary haematopoiesis). The splenic architecture is maintained, with evident lymphoid follicles. There are multifocal increased numbers of hemosiderin macrophages. |

Of all the aforementioned lesions, and consistent with prior literature^44^, splenic enlargement based on absolute spleen weight was noted in a subgroup of aged mice **(Fig. S2A)**. However, the splenic weight index (SWI) did not reveal a significant increase **(Fig. S2B)**, indicating that splenic enlargement either accompanies or occurs at a lower rate compared to body weight gain with aging. Moreover, spleens from Old-aged mice presented with extramedullary hematopoiesis, characterized by red pulp megakaryocytes with hemosiderosis **(Figs. S1F-F’)**. To determine the relative contribution of individual splenic compartments to the overall enlargement, we developed two independent morphometric methods, an HE-based and an IF-based, both calculating the White-to-Red-Pulp Ratio (WRPR) **(Fig. S2C)**. Both methods correlated well with each other, and revealed a trend for WRPR decline with aging, but neither showed statistical significance across the age groups **(Figs. S2D-F)**. In addition, we assessed the white and red pulp fractions independently, but again, neither method revealed any changes with aging **(Figs. S2G-L)**. In conclusion, splenic enlargement is an incidental finding in only a subset of Old mice, and seems to occur due to the concerted increase of both white and red pulp.

We then macroscopically assessed the extracted thymi and found a significant decline of thymic weight with aging **(Fig. 1C)**. In consistency, the thymic weight index (TWI) continuously declined from a mean of ∼3.46 at the Young-Adult to a mean of <0.7 at the Old group **(Fig.1D)**. This rapid decline in thymic size **(Figs.1C-D)** is well-characterized in mice^45^, and is consistent with thymic decline in humans after reaching puberty^46–48^. In conclusion, a macroscopic analysis of the thymus in female mice reveals evidence of age-dependent thymic involution^45–48^, and as such, it represents an efficient *in vivo* model for the development of microanatomical and ultrastructural morphometries, using our digital pathology framework.

### Microscopic Assessment of Cortex-Medulla Compartmentalization

Our subsequent studies focused on establishing morphometric assessments of the thymic structure, to draw functional correlates of thymic function and thymopoietic capacity. Based on International Harmonization of Nomenclature and Diagnostic Criteria (INHAND) histopathology guidelines by multiple Societies of Toxicological Pathology^27,28,49^, the volumetric ratio of thymic cortex to medulla (i.e., corticomedullary ratio, CMR) is the strongest indicator of functional status, and may be altered in pathology and/or pharmacologic interventions, such as in endogenous cortisol production in response to chronic stress, or following exogenous glucocorticoid or cyclosporin administration^1,27,28,49,50^. The guidelines provided by INHAND, however, are based on evaluating the ratio subjectively, thus leading to operator-dependent biases^27^. Based on the guidelines^27^, here, we sought to standardize the “corticomedullary ratio” as a reproducible and measurable index, to inform on the structural and functional status of the thymic parenchyma.

Our pipeline involves three independent methods that calculate corticomedullary ratio in a digital pathology framework **(Figs. 2A-A’’’)**. Thymus sections **(Fig. 2A)** are first stained either with a Keratin-5/Keratin-8 (KRT5/8) dual immunofluorescence staining protocol (Slide 1), or with HE (Slide 2), and digital whole slide-scanning is used to generate whole tissue sectioned images. The cortical and medullary compartments are then digitally demarcated as individual regions of interest (ROIs) in either the KRT5/8 or the HE modality **(Fig. 2A’)**. In the HE modality, the cortex appears darker, because cortical thymocytes are smaller in size and tightly arranged in their physical space, when compared to their medullary counterparts, thus generating higher density of hematoxylin^+^ nuclei **(Fig. S3A)**. In the IF modality contrariwise, the specific keratin filament immunolabeling signifies topology of specific thymic epithelial cell (TEC) compartments, with KRT5 exclusively labeling medullary TEC (mTEC), whereas KRT8 labeling all cortical TEC (cTEC), along with a prominent subset of mTEC **(Figs. S3B-B’’)**^51–54^. The cortical parenchyma is thus defined as KRT5^-^KRT8^+^, while the medullary parenchyma as KRT5^+^KRT8^+^ **(Figs. S3B’-B’’)**. Besides the positivity/negativity in the keratin signal, the nature of the cTEC/mTEC networks is also distinctly diverse, with cTEC appearing with “meshwork” architecture while mTEC densely distributed as isolated cellular entities **(Figs. S3B-B”)**. In either modality (HE or KRT5/8 IF), a clear distinction of the corticomedullary junction (CMJ) is demonstrated **(Figs. S3A-B)**, and is used for the digital demarcation of the non-overlapping cortical and medullary ROIs **(Fig. 2A’’)**. CMR is then defined as the ratio of surface areas of the two thymic compartments, and abbreviated as CMR*_WT-HE_* (for the HE modality), and CMR*_WT-IF_* (for the IF modality) **(Fig. 2A’’’ and equation #1)**.

**Figure 2.**
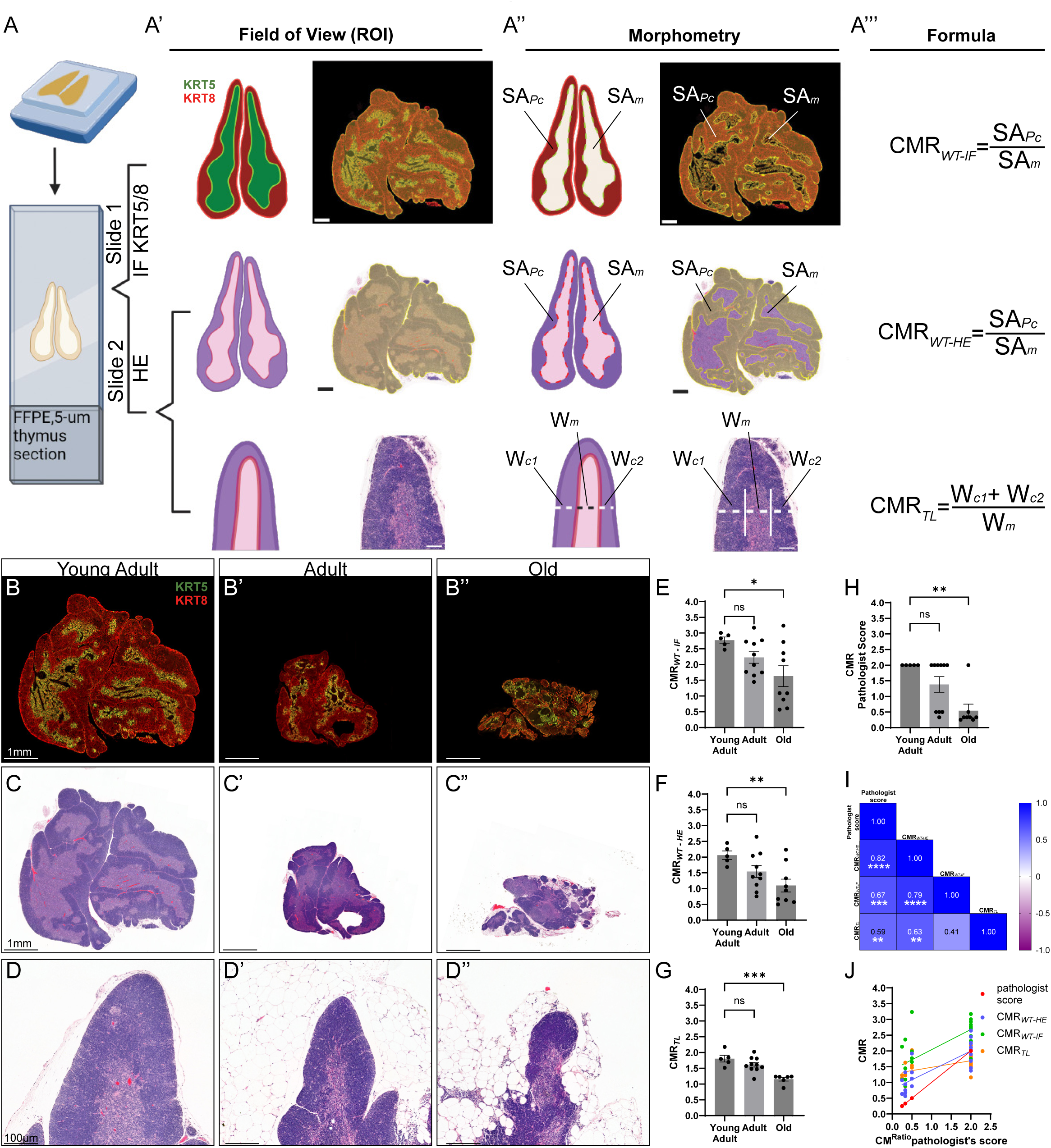
Morphometric assessment of corticomedullary ratio. **(A-A’’’)** Digital pathology pipeline for the “whole-thymus” (1^st^ and 2^nd^ row) and “thymic-lobule” (3^rd^ row) methods, using Keratin-5/8 (KRT5/8) IF (1^st^ row) and HE (2^nd^ and 3^rd^ row) staining. From left to right: 5µm thick sections from FFPE blocks are HE- and IF-stained (A). Representative fields-of-view (FOVs) are then selected as annotations using digital software like Caseviewer; whole thymi (yellow hue) are selected for the “whole-thymus” approaches, whereas single thymic lobules are selected for the “thymic-lobule” approach (A’). Following the annotations, representative regions-of-interest (ROIs) are created; the cortical and medullary parenchyma are independently generated for the “whole-thymus” approaches, whereas one-dimensional line ROIs segment the thymic lobule into cortical and parenchymal segments (A’’). Following ROI generation, formulas are applied for calculating the corticomedullary ratios (CMR); for the “whole-thymus” approaches the CMR*_WT_* takes into consideration the surface areas (SA) of the ROIs, whereas for the “thymic-lobule” approaches, the CMR_TL_ takes into consideration the widths (W) of the ROIs (A’’’). *Abbreviations: SA_Pc_, surface area of cortical parenchyma; SA_Pm_, surface area of medullary parenchyma; CMR, corticomedullary ratio; W_c1_, width of cortical segment 1; W_c2_, width of cortical segment 2; W_m_, width of medullary segment.* **(B-D)** Representative images of thymi across the three age groups, for CMR assessments. From top to bottom: Whole slide IF images of thymi, representative of Young-Adult (B), Adult (B’) and Old (B’’) mice; Whole slide HE images of thymi, representative of Young-Adult (C), Adult (C’) and Old (C’’) mice; HE thymic lobule images of thymi, representative of Young-Adult (D), Adult (D’) and Old (D’’) mice. *Note: the thymus depicted in (B-C) is also shown for demonstration purposes in the pipeline (A).* **(E-H)** Quantification of corticomedullary ratio (CMR), using digital (E-G) and traditional pathology approaches (H). Shown in sequence: “whole-thymus” approach using IF staining (CMR*_WT-IF_*) (E); “whole-thymus” approach using HE staining (CMR*_WT-HE_*) (F); “thymic-lobule” approach using HE staining (CMR*_TL_*) (G); blinded pathologist’s scoring, using an arbitrary scale (H). *One-Way ANOVA; *p*≤*0.05; **p*≤*0.01; ***p*≤*0.001; ****p*≤*0.0001; ns, not-significant.* **(I-J)** Analysis of linear correlation among the different established methods of CMR assessment. The correlations are assessed using Spearman’s correlation coefficient analysis, and visualized with a Spearman’s correlation matrix showing the corresponding Spearman rho and p-values for each pair (I), as well as with a scatter plot with simple linear regression line between the pathologist’s score and each of the newly established methods (J).

The whole-thymus approaches described above may sample a large area for analysis, but they are more prone to errors regarding the orientation of the organ’s axis in the paraffin block, as well as to the depth of block “trimming” prior to sectioning (e.g., superficial trimming would result in biased cortex domination due to the tangential cutting of the lobules). To circumvent these issues, we developed another independent method, by standardizing the quantified ROI across different samples. The method exclusively relies on images of individual thymic lobules, whose corticomedullary junction orientation is arranged in parallel fashion across the two sides of the medullary parenchyma. In these lobules, a one-dimensional line is then drawn in perpendicular fashion to the orientation of the corticomedullary junction **(Fig. S3C)**. The cortical and medullary segment widths are then measured on this vector **(Fig. S3C)**, and the ratio of absolute widths is calculated for each mouse **(Fig. 2A’’’ and equation 2)**. Due to the measurement of corticomedullary ratios using standardized microanatomical features, i.e., thymic lobules with a predetermined orientation, this method (CMR*_TL_*) is more consistent among replicate experiments. It could be limited, however, by pathologies resulting in loss of thymic lobulation, etc.

An in-depth analysis of functional adaptations of thymic compartmentalization during aging has previously demonstrated the numeric decline of cTEC, leading to a reduced thymopoietic potential of the aged thymus^55^. Interestingly these data have further documented that mTEC do not seem to decline as fast with aging, although many mTEC subsets tend to lose their terminal differentiation and present with a progenitor-like phenotype^55^. These observations suggest an expected reduction of the corticomedullary ratio in aged mice^28^. To confirm this hypothesis, we assessed CMR using all three methods described, and successfully captured its expected decrease with aging **(Figs. 2B-G)**. Although the tendency of decline was clear after adolescence, none of the three methods captured any difference (p>0.05) between Young-Adult and Adult groups **(Figs. 2E-G)**. However, there was significant reduction between Young-Adult and Old groups for all three methods **(Figs. 2E-G)**, suggesting that our morphometries can efficiently capture the expected disturbances in thymic compartmentalization with older age.

To determine a potential deviation of our proposed morphometries from the accepted and established methods of corticomedullary ratio calculation, we performed a subjective analysis of corticomedullary ratio, based on the INHAND guidelines^26–28^. The measurements were made by a board-certified veterinary pathologist, who was “blinded” to the mouse age-groups, using an arbitrary scale of categorical CMR values. This method also successfully captured the expected decrease of the corticomedullary ratio with aging **(Fig. 2H)**. Importantly, the pathologist’s scores correlated best with the CMR*_WT-HE_*, when compared to the CMR*_WT-IF_* and CMR*_TL_* approaches **(Figs. 2I-J and S3D-F)**, which was expected given that the blinded pathologist also utilized whole-thymus HE slides, to perform quantifications. When the three newly-developed methods were compared to one another independently, we found that the best correlation was between whole-thymus approaches (i.e., CMR*_WT-HE_* and CMR*_WT-IF_*) with spearman rho of 0.79 (p<0.001). As expected, the worst correlations were observed between the CMR*_TL_* and other approaches **(Figs. 2I and S3G-I)**. Despite that the CMR*_TL_* deviated numerically from all other methods, it was the method that presented with the least variability (i.e., standard deviation) for each mouse age-group **(Fig. 2G)**, an expected finding, given that it is exclusively based on a pre-defined microanatomical unit. These observations demonstrate that the proposed methods exhibit high orthogonality and could be used complementarily to evaluate thymic compartmentalization.

### Topological Assessment of the Cortical Thymic Epithelial Cell (cTEC) Network

In post-embryonic thymus, cTEC develop with meshwork architecture, and their functional maturation requires thymocyte signals beyond the DN1 stage^2,56^ . Reciprocally, the cTEC are incremental for properly educating thymocytes and for their progressive maturation through most developmental stages^2,4,56,57^ . The meshwork architecture of the cTEC population is defiled in certain thymic pathologies, including the natural age-dependent thymic decline, as well as acute thymic involution induced by cytotoxic stressors^7,24,26–28,58^. As the next step of our digital pathology pipeline, we therefore propose a rationalized approach to establish quantifiable features of cTEC meshwork architecture, to characterize its functional thymopoietic capacity.

The first feature, herewith termed “cTEC Volume Index” (VI*_cTEC_*), reports the cTEC volume as a fraction of the parenchymal mass of the cortex. The analysis involves an initial binarization of the KRT8^+^ signal based on intensity thresholding to generate a “cTEC mask”, then reporting the KRT8^+^ surface area as a proportion of the total surface area of the cortical parenchyma **(Figs 3A-C’, 3F and equation 3)**. Because there are no other cells known to express KRT8 in the thymic cortex apart from the cTEC stroma^54^, an increased ratio would thus indicate cTEC hyperplasia or hypertrophy (i.e., increased cTEC volume per unit of thymic parenchyma), while a decreased ratio would indicate cTEC atrophy or hypoplasia (i.e., decreased cTEC volume supporting thymopoiesis) **(Fig. 3G)**. It should be noted though that the dual KRT5/KRT8 immunofluorescence is critical for the assessment of this feature, because KRT5^+^KRT8^+^ regions correspond to the medulla, and thus should be excluded from the analysis **(Figs. 3A-A’, S3B-B’’)**.

**Figure 3.**
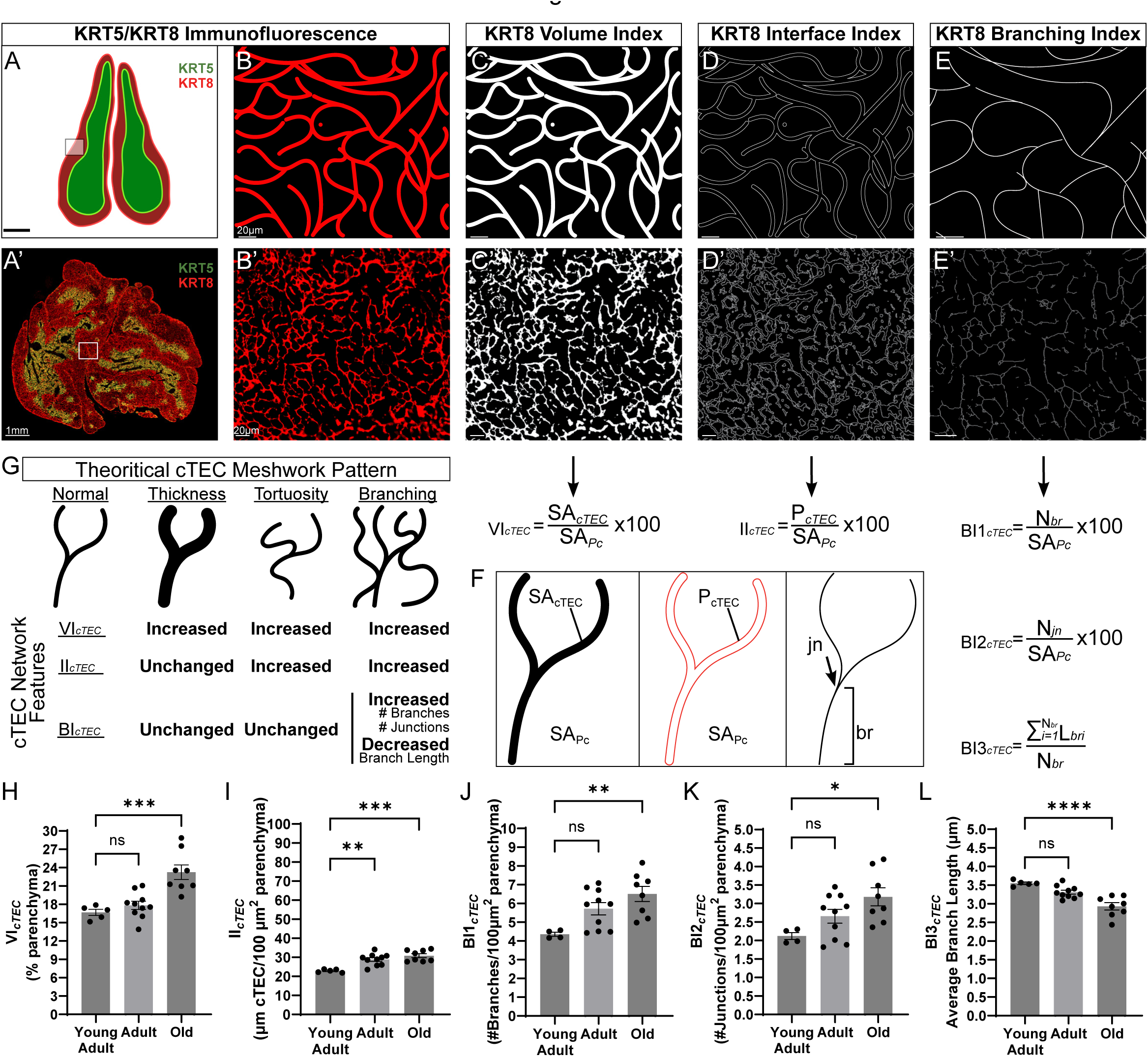
Topological assessment of the cTEC network. **(A-F)** Digital pathology pipeline for the topological assessment of cTEC meshwork architecture. **(A-A’)** Whole thymi are stained for KRT5/8 immunofluorescence, and cortical regions are identified as KRT5^-^KRT8^+^. **(B-B’)** Thymic cortex field-of-view (FOV), depicting meshwork architecture of the cTEC network. **(C-C’)** The binarization of the image shown in (B) via KRT8 signal intensity thresholding, creates a “mask” of the cTEC network. Accompanying formula indicated with the arrow. **(D-D’)** Development of outline from the binary mask shown in (C), revealing the interface area of the cTEC network. Accompanying formula indicated with the arrow. **(E-E’)** Skeletonization of the binary mask shown in (C), revealing the branching characteristics of the cTEC meshwork. Accompanying formulas indicated with the arrow. **(F)** Illustration of a cTEC network part that can be used as “key” for the formulas shown in (C-E). *Abbreviations: VI_cTEC_, cTEC volume index; II_cTEC_, cTEC interface index; BI_cTEC_, cTEC branching index; SA_cTEC_, cTEC surface area; SA_Pc_, surface area of cortical parenchyma; P_cTEC_, cTEC perimeter; jn, junction; br, branch.* (G) Theoretical adaptations of the cTEC network topology, along with expected changes in cTEC network morphometries following these adaptations. **(H-L)** Morphometric assessment of cTEC network topology, using cTEC volume index (VI*_cTEC_*) (H), cTEC interface index (II*_cTEC_*) (I); cTEC branching indices, i.e., BI1*_cTEC_* (J), BI2*_cTEC_* (K), and BI3*_cTEC_* (L). *One-Way ANOVA; *p*≤*0.05; **p*≤*0.01; ***p*≤*0.001; ****p*≤*0.0001; ns, not-significant*.

The second feature, herewith termed “cTEC Interface Index” (II*_cTEC_*), informs on the surface area of the cTEC meshwork, readily exposed for direct cTEC-thymocyte interactions. As the developing thymocytes travel through the cortex, they are juxtaposed and continuously come in contact with MHC I/II-self-peptide complexes that are presented in the surface of cTEC^4,59–61^. To draw functional correlates of this process, we implemented an extension of the previous analysis that generates the cTEC mask based on KRT8^+^ signal intensity thresholding. Besides calculating the VI*_cTEC_* from this binary mask, an “outline” of the mask is generated **(Figs. 3D-D’)**, and its perimeter is measured and normalized to a given surface area of cortical parenchyma **(Figs. 3F and equation 4)**. In case of VI*_cTEC_* increase for instance, a corresponding II*_cTEC_* would inform on whether the observed cTEC volume increase is due to increase in cTEC cell body thickness, or due to increase of cTEC network tortuosity, branching morphogenesis, etc. **(Fig. 3G)**.

The third feature, herewith termed “cTEC Branching Index” (BI*_cTEC_*), informs on more detailed topological and structural descriptors of the cTEC network. Because cTEC exhibit dendritic morphology, they create a vast meshwork, characterized by establishment of cytoplasmic junctions (nodes) and branches **(Figs. S3B-B’)**. As above, this feature also considers a cTEC mask, as generated to calculate VI*_cTEC_* and II*_cTEC_*, but at higher magnification. The cTEC mask is “skeletonized” using an ImageJ plugin^38^, an algorithm that reduces the binary cTEC mask to one-pixel thickness but retaining the original object representation **(Figs. 3E-E’)**. This transformation allows for straightforward quantification of network descriptors, such as number of branches and junctions, both normalized to a given surface area of cortical. With this method, the “average branch length” of all branches in the cTEC skeleton is also automatically calculated, when the network skeleton is analyzed **(Figs. 3E-E’, 3F and equations 5-7)**. With regards to interpretation, the three BI*_cTEC_* features, i.e., BI1*_cTEC_*, BI2*_cTEC_*, and BI3*_cTEC_*, collectively illustrate the nature of the cTEC meshwork complexity, specifically its tortuosity and branching capacity, and offer complementary information when assessed along with other descriptors, such as VI*_cTEC_* and II*_cTEC_* **(Fig. 3G)**.

A previous study has performed quantitative assessments of age-dependent lesions in cTEC and mTEC, and reported structural and functional deficits of the former compartment in the murine thymus^62^. In agreement, our results indicate a significant increase (p<0.001) of VI*_cTEC_* in older mice, compared to younger mice **(Fig. 3H)**. However, this cTEC volume “increase” should be cautiously interpreted in the context of overall numerical cTEC and thymocyte decline, known to occur with aging^63^. The cTEC network is generated through long, looping structures forming an intracellular labyrinth of multiple “voids” in each TEC, inside which groups of 2-10 cortical thymocytes reside^62^. As such, a reduced number of thymocytes due to aging could lead to the “narrowing” of these voids, and the artificial resemblance of the cTEC network as more “massive” in nature. Along these lines, the remaining network descriptors, e.g., II*_cTEC_*, BI1*_cTEC_*, and BI2*_cTEC_* also revealed a significant increase in older mice **(Figs. 3I-K)**, demonstrating the dynamic shift of cTEC/thymocyte mass ratio. In consistency with these data for example, the relative area occupied by CXCL12^+^ cells, a homing chemokine exclusively produced by cTEC, was found to be larger in older thymi in humans, again due to the reduction of thymus size with aging^64^.

When we specifically evaluated the branching capacity of the cTEC network in older mice, we found significant increase (p<0.05) in the relative number of branches and junctions, when compared to the younger mice **(Figs. 3J-K)**. This unorthodox observation could be explained by the hypertrophic growth of the few surviving TEC in older thymi, and/or the simultaneous cTEC network collapse, due to the sparse distribution of resident thymocyte populations within the cTEC labyrinthine voids. One study has recently reported inconspicuous development of cTEC projections in old mice^62^, thus challenging our observations. In consistency with our results however, other studies have reported cTEC hypertrophy as a result of Lymphotoxin-β receptor (LTβR) reduced expression or deficiency^65^, which is normally observed during aging in lymphoid organs^66,67^. In accordance with our finding showing higher number of branches and junctions, we also found significant decrease in our final network descriptor, the average branch length (BI3*_cTEC_*), in the Old mouse group **(Fig. 3L)**. This finding is highly consistent with the hypertrophic nature of the surviving cTEC at old age, because newly formed branches typically have less time to grow, and are thus expected to be shorter **(Fig. 3F)**.

In summary, a detailed analysis of cTEC network topology has demonstrated a substantial remodeling towards hypertrophic growth with aging. cTEC hypertrophy in this case, however, is not characterized by an abnormal thickening of individual cTEC branches, but by the *de novo* branching morphogenesis and increased tortuosity of the network. These findings collectively suggest that our newly proposed morphometries must be used non-redundantly, to describe the precise topology of the cTEC network across thymus health and disease.

### Ultrastructural Topography of the cTEC Network

To validate our findings and gain further resolution on cTEC network remodeling with aging, as visualized through the morphometric assessments using immunofluorescence **(Fig. 3)**, we combined scanning electron microscopy of OTO-sections (OTO-SEM) with transmission electron microscopy (TEM) in mice undergoing age-dependent thymic involution. The cellular component of the cTEC meshwork, also known as “cytoreticulum”, is identified in both OTO-SEM and TEM by the characteristic presence of cytoplasmic tonofilaments, vacuoles, and the occasional presence of desmosomes between interconnecting cTEC projections **(Fig. S4)**, as reported^68–72^. These distinguishing epithelial features provide validation of cTEC identity, especially when they are projected in proximity to cTEC nuclei, or when the cytoplasmic projection is shown to bud off the cTEC on the same z-plane **(Fig. S4C; asterisk)**. As opposed to thymocyte nuclei, which often contain conspicuous heterochromatin patches giving a “rosette- or flower-shaped” impression, the cTEC nuclei are more euchromatic and easily distinguished **(Figs. S4C-D)**. However, the identification of such epithelial features becomes more crucial when tracking trajectories of cytoreticulum segments among developing thymocytes, either because the cTEC nucleus is farther away from the respective field-of-view, or exists in a different z plane **(Fig. S4D; asterisk)**.

Because the cTEC network develops meshwork architecture, single section analyses would provide valuable, albeit limited information. We thus obtained 3-dimensional reconstructions at high resolution using 3D OTO-SEM Array Tomography^73^. Backscattered OTO-SEM revealed the ultrastructural details of cTEC cytoreticulum, while it partially or completely enveloped the thymocytes under various stages of their development in the outer cortex **(Figs. 4A-B)**. The cytoreticulum (with/without inclusion of the cTEC nucleus or nuclei) was then outlined in each z-stack independently, and reconstructed **(Figs. 4A-B)**. 3D reconstruction views of the cTEC meshwork confirmed that the cytoreticulum formed well-circumscribed labyrinthine “voids”^62^, each homing groups of ∼1-10 developing thymocytes **(Figs. 4A-B)**. However, viewing the cytoreticulum in single OTO-SEM sections fails to demonstrate the continuity of this unique architectural pattern **(Figs. 4C-E)**, as evidenced by the 3D reconstruction **(Figs. 4A-B)**. Thymocytes within these voids are in partial or complete contact with the adjacent cytoreticulum **(Figs. 4A-E)**, suggesting that the cytoreticulum reflects to regions rich in MHC I/II-peptide complexes directly presented to enveloped thymocytes. At first glance, one could erroneously assume that cTEC-thymocyte interactions improve with older age, given the numerical decline of thymocytes and the concurrent cTEC hypertrophy **(Figs. 3 and 4A-E)**; in other words, fewer thymocytes now correspond to larger cTEC volumes. However, single section analyses using TEM reveals that cortical thymocytes tend to lose their strong adherence to the nearby cTEC meshwork, thus creating more profound “empty spaces” around thymocytes with aging **(Figs. 4F-H; arrows)**. These data collectively suggest that the ability of thymocytes to adhere to the cTEC network within the labyrinthine voids, is corrupted in aged mice.

**Figure 4.**
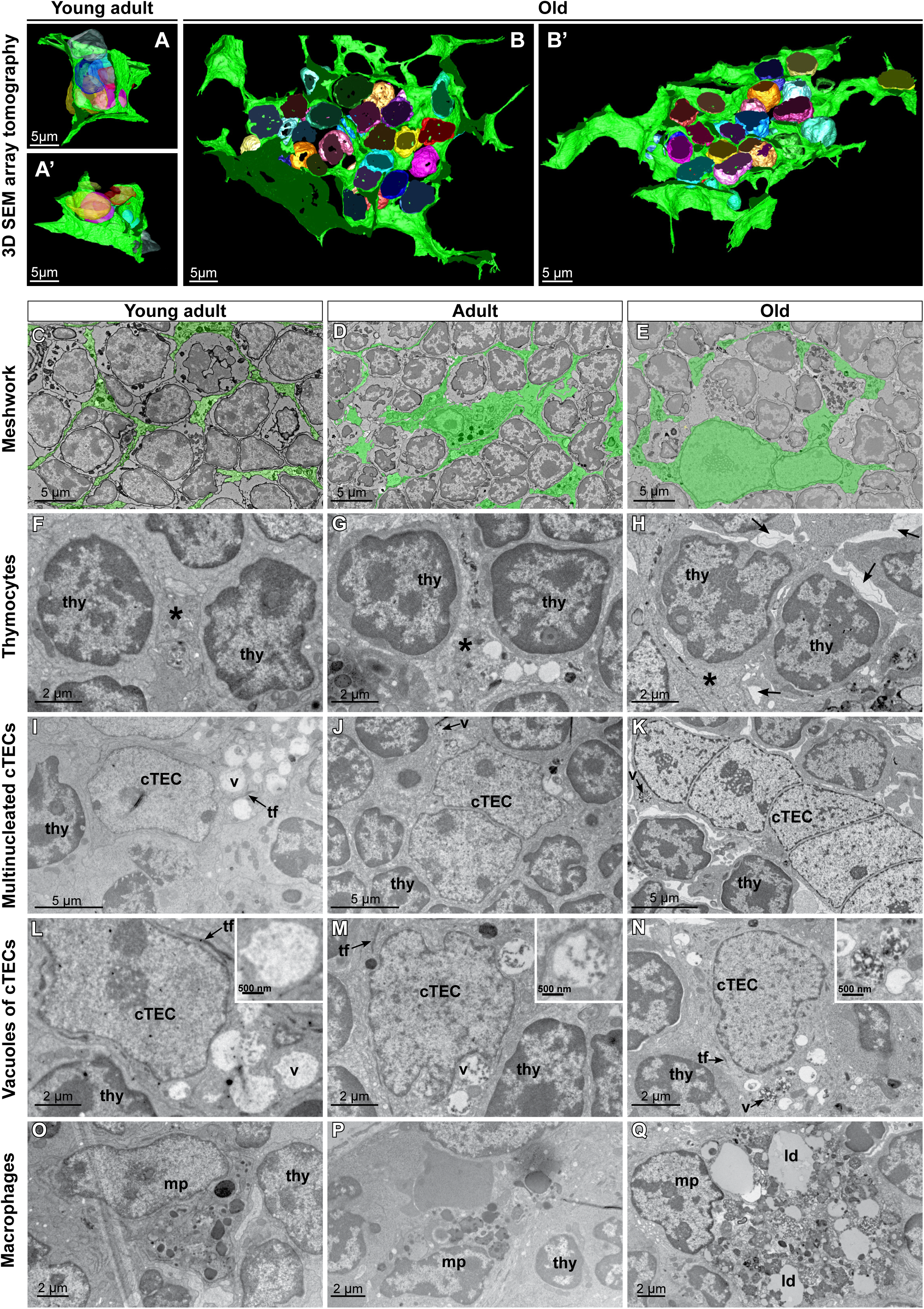
Ultrastructural topography of the cortical parenchyma and the cTEC meshwork. **(A-B)** Three-dimensional reconstruction of the cTEC network, shown in green, with translucent multicolored nuclei of developing thymocytes shown surrounded by the network in one Young Adult (A-A’) and one Old (B-B’) mouse. A’-B’ show different angles/representations of the same image. **(C-E)** Ultrastructural architecture of cTEC network (green overlay) creating labyrinthine voids and enveloping developing thymocytes in Young-Adult (C), Adult (D), and Old (E) mice, using OTO-SEM). **(F-H)** Ultrastructural assessment of cTEC-thymocyte interactions in Young-Adult (F), Adult (G), and Old (H) mice, showing an increase of empty spaces in-between in older mice. *thy, thymocyte; asterisk, cTEC cytoplasmic projections; arrows, empty spaces among cTEC and thymocytes.* **(I-K)** Ultrastructural assessment of cTEC multinucleation in Young-Adult (I), Adult (J), and Old (K) mice, showing evidence multinucleation with aging. *thy, thymocyte; v, vesicle; tf, tonofilament.* **(L-N)** Ultrastructural assessment of vacuoles in the perinuclear cytoplasm of cTEC, in Young-Adult (L), Adult (M), and Old (N) mice, showing increased electron density with aging. Magnified inserts show the vesicles labeled as “v” in the corresponding images. *thy, thymocyte; v, vesicle; tf, tonofilament.* **(O-Q)** Ultrastructural assessment of cortical macrophages in Young-Adult (O), Adult (P), and Old (Q) mice, showing progressive deterioration with aging, characterized by increased size and accumulation of lipid droplets. *mp, macrophage; ld, lipid droplet*.

Evidence of cTEC cytoreticulum surrounding small groups of thymocytes was confirmed in both Young-Adult **(Fig. 4A)** and Old mice **(Fig. 4B)**. In Young-Adult mice, however, cTEC were significantly smaller in size, each creating 1-2 labyrinthine voids surrounding a substantial number (∼10) of lymphocytes each **(Fig. 4A)**. On the contrary, cTEC were more massive in Old mice, each creating a large number of labyrinthine voids but surrounding a significantly smaller number of thymocytes each, i.e., 1-2 thymocytes **(Fig. 4B)**. These data are consistent with our novel premise that age-dependent hypertrophy of the cytoreticulum represents a compensatory mechanism to the numerical decline of its population, in an effort to preserve the thymopoietic capacity of the cortical parenchyma.

An ultrastructural feature that could further support the establishment of cTEC hypertrophy during aging as a means to compensate for the numerical decline of the cTEC population, is the emergence of multinucleated cTEC **(Fig. 4I-K)**. In general, multinucleation in thymic epithelial cells has been observed in involuted thymi using electron microscopy^74^, confirming the legitimacy of our observations. It is uncertain whether such multinucleated cTEC develop via cTEC fusion or via a pathological mechanism that includes disturbed cell cycle control. However, these cTEC appear non-apoptotic **(Fig. 4I-K)**, implying they might imitate other cells in the body adopting syncytial configuration, in response to increased metabolic demands^75^. Indeed, ∼24-month-old mice exhibited cTEC with up to 5-6 nuclei arranged in a “caterpillar” pattern **(Fig. 4K)**. Even though we identified a few bi-nucleated cTEC in younger mice, these findings have been rare instances and not consistent. These observations could formulate the working hypothesis that the rate of cTEC multinucleation in the cortex is directly proportional with age, although a statistical conclusion may not be easily drawn from qualitative TEM data, due to the low yield of observable events. As a newly proposed feature in our pipeline, however, cTEC multinucleation could represent a key remodeling event observed in various thymic pathologies, especially those associated with underlying involution.

Cytoplasmic vacuoles in TEC had been observed in electron microscopy studies, but their precise functional roles were not appreciated at that time^69,76^. It is now established that cTEC employ both thymoproteasome-dependent and macroautophagy-dependent mechanisms to digest, produce, load, and present self-peptides on their surface via the major histocompatibility complex (MHC)^77,78^. These degradation pathways are tightly controlled in the normal thymus, to prevent an unrestrained disturbance of TEC function and homeostasis. Ultrastructural analysis of cTEC vacuoles defined their identity as auto(phago)lysosomes **(Figs. S4A-A’)**, implicating them in the macroautophagy machinery^79^. We found that cTEC autolysosomes are distributed throughout the cTEC body, including the perinuclear cytoplasm, as well as more distally within the cytoplasmic projections **(Figs. 4L-N and S4)**. It is very possible that these vacuoles are strategically positioned near the sites of self-peptide loading into MHC molecules. In Young-Adult mice, autolysosomes appear as single-membrane vesicles containing mostly amorphous (partially-digested) material in their lumen, resembling either a string of pearls, or slightly conspicuous vesicular/membranous inclusions **(Figs. 4I&L)**. Also, a vast portion (>80%) of their content is electron-lucent **(Fig. 4L)**. During aging, however, another phenotype of autolysosomes is encountered. These are packed with a higher proportion (∼60-80%) of amorphous electron-dense material **(Figs. 4M-N)**, indicative of entire organelle inclusion and destruction. Based on prior knowledge, these electron-dense autolysosomes are suggestive of pathological stress-induced macroautophagy^80^, occurring during acute or chronic thymic involution. Here, by focusing on the measurement in a specific ultrastructural compartment, i.e., the perinuclear cytoplasm, we propose that morphological assessment of electron density of cTEC vacuoles draws functional correlates on the cTEC ability to drive thymopoiesis and adaptability to stress.

In general, cortical macrophages exert multiple functions with the most exemplified ones comprising phagocytosis of thymocytes that fail positive selection, and maintenance of blood-thymus barrier, i.e., scavenging of foreign antigens that leak into the thymic parenchyma^81^. Due to their increased phagocytic activity, macrophages are often deemed the “tingible-body” or “starry sky” pattern description in routine histology^27^. Varying degrees of macrophage activity was observed in mice across the different age groups, with the most prominent feature the presence of secondary lysosomes in their cytoplasm **(Figs. 4O-Q)**. With aging, however, lipid-laden macrophages (LLM) were clearly evident **(Fig. 4Q)**, and associated with lipid droplets of various sizes and densities in their cytoplasm, as reported^82^. These data confirm previous findings that cortical macrophages undergo age-dependent attrition, as opposed to medullary counterparts that persist through life^81^. In general, LLMs have been associated with disease progression in inflammatory conditions^83^, and LLM indexes have been successfully used as diagnostic tools^84^. LLM presence in the cortex, as assessed by our TEM analysis, could thus be explored for its potential to inform on the functional characterization and disease progression in the thymus.

Together, the combined OTO-SEM and TEM studies detailed here, complement the digital pathology pipeline discussed in the previous sections. By offering ultrastructural insights into the cTEC meshwork remodeling with aging, these studies not only validate established knowledge in the field, but also enhance our understanding of thymic structure and function under various thymic pathologies, illustrating the intricate changes at the ultrastructural level.

### Morphometric Analysis of the Medullary Perivascular Space (PVS)

After successful negative selection, the mature SP4/8 thymocytes egress from the medullary parenchyma via tightly controlled chemokine signals, contextualized in microanatomical regions, called perivascular spaces (PVS). Medullary vessels that are specialized for thymocyte egress have been demonstrated to contain double membranes, an inner boundary consisting of pericytes wrapped around blood vessel endothelial cells and an outer boundary consisting of adventitial cells of mesenchymal/fibroblast origin^17,39,85–89^. The two boundaries simultaneously circumnavigate the medullary blood vessel and encompass a well-demarcated area that excludes mTEC, thus appearing as an epithelium-free region^39^ **(Fig. 5A)**. SP4/8 thymocyte egress is dependent upon chemokine signals, most notably sphingosine-1-phosphate (S1P), whose controlled biogenesis/degradation, orchestrates their stepwise emigration, first from the medulla into the PVS and then, from the PVS into the blood vessel^90–92^. Because many natural or pathological conditions affect the PVS and thymic output^93–96^, the composition and size of the PVS constitutes an important topological feature reflecting to the ability of the medulla to pump out mature thymocytes to the periphery.

**Figure 5.**
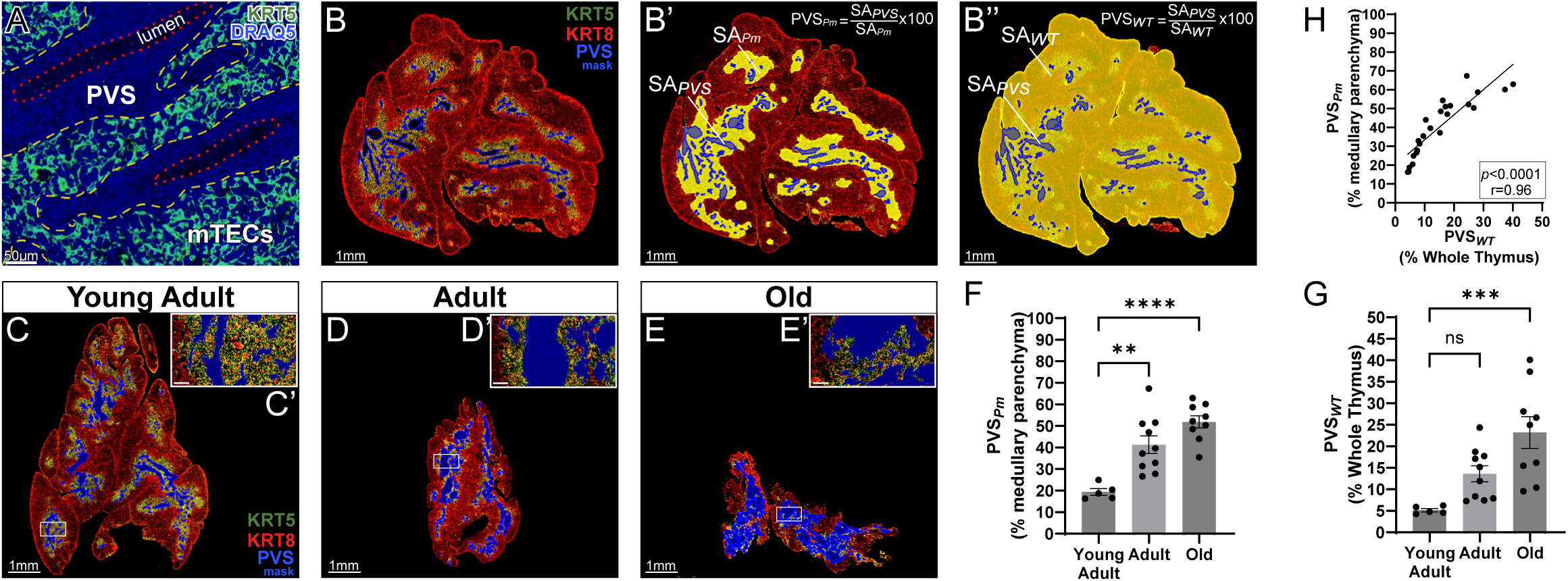
Morphometric assessment of the medullary perivascular space. **(A)** Representative image of thymic medulla at high magnification, highlighting the mTEC (KRT5^+^) network and the epithelium-free areas, corresponding to the perivascular spaces (PVS). A special blood vessel stain has not been performed, but blood vessel lumens are apparent in these sections, based on absence of cell nuclei (DRAQ5). *Yellow-dashed line, outer boundary of the PVS; red-dotted line, inner boundary of the PVS, corresponding to the blood vessel wall.* **(B-B’’)** Digital pathology pipeline for assessing the fraction of the PVS within the thymus. KRT5/8 IF staining reveals the cortical (KRT5^-^KRT8^+^) parenchyma, the epithelium-containing medullary (KRT5^+^KRT8^+^) parenchyma, and the medullary PVS (KRT5^-^KRT8^-^), here depicted in blue mask (B’). Morphometric assessment of the PVS as a fraction of the medulla-only (B’), or the entire thymus (B’’). The regions-of-interest used for the normalization, i.e., medulla only, and whole thymus, are demarcated with yellow color in (B’) and (B’’), respectively. The formulas used to calculate the two versions of PVS fraction are shown in the upper right corner. *Abbreviations: SA_PVS_, surface area of the PVS; SA_Pm_, surface area of medullary parenchyma; SA_WT_, surface area of whole thymus.* **(C-E)** Representative images of thymi along with magnified inserts (C’-E’) across the three age groups, for morphometric assessment of the PVS. Shown from left-to-right: Young-Adult (C-C’), Adult (D-D’), and Old (E-E’) groups. **(F-G)** Quantification of PVS fraction, using digital pathology approaches. Shown in sequence: PVS as fraction of the medulla (PVS*_Pm_*) (F), and PVS as fraction of the whole thymus (PVS*_WT_*) (G). *One-Way ANOVA; *p*≤*0.05; **p*≤*0.01; ***p*≤*0.001; ****p*≤*0.0001; ns, not-significant.* **(H)** Analysis of linear correlation between the two established morphometric methods. The correlations are assessed using Spearman’s correlation coefficient analysis with Spearman rho and p-values, and visualized with a scatterplot with simple linear regression line between the two methods.

Previous studies have reported the quantification of the PVS fraction in the parenchyma of the thymus, either in a non-compartmentalized (i.e., in relation to the surface area of the whole thymus), or a compartmentalized (i.e., as a fraction of medullary parenchyma only) fashion^12,94^. The two strategies are informative when assessing either the medullary functions specifically, or in respect to the total thymopoietic capacity, respectively. Based on prior knowledge that the medullary PVS is an epithelium-free area^17^, we drew regions of interest (ROIs) devoid of the KRT5^+^KRT8^+^ epithelium around the medullary PVS **(Figs. 5A-B)**, and calculated the proportion of the surface area covered by the demarcated PVS within the KRT5^+^KRT8^+^ surface area of the medullary region (PVS*_Pm_*) **(Fig. 5B’ and equation 8)**, or within the KRT8^+^ (whole-thymus) surface area (PVS*_WT_*) **(Fig. 5B’’ and equation 9)**. In consistency with prior observations in human thymi^94,97,98^, we found that the PVS fraction is significantly increased over time, regardless of the morphometry used **(Figs. 5C-G and S5A-C)**. Impressively, approximately 50% of the medullary parenchyma and 25% of the entire thymic parenchyma corresponds to the PVS zone in the Old group **(Figs. 5F-G)**. In addition, there is a significant linear correlation (Spearman rho = 0.96) between the two morphometries **(Fig. 5H)**, indicating that the methods are in fact orthogonal, and as such, equally suitable for studying factors affecting thymocyte emigration.

### Topological Assessment of the Medullary Thymic Epithelial Cell (mTEC) Network

As opposed to cTEC, mTEC are scattered in freeform arrangement without developing any meshwork architecture in the medulla^4^. The interaction among different mTEC subsets and the single-positive (SP) thymocytes, controls the fate and success of negative selection, and eventually thymocyte egress^4^. Due to the absence of meshwork architecture, an identical pipeline to that of cTEC, e.g., including skeleton analysis, could not be implemented for mTEC topology. However, as in the case of cTEC, SP4/8 thymocytes also interact through direct physical contact with various mTEC subsets to complete their developmental journey^4^. As such, the two features, “mTEC Volume Index” (VI*_mTEC_*) and “mTEC Interface Index” (II*_mTEC_*), were employed in a similar fashion, because they are equally informative with the cTEC equivalents **(Figs. 6A-D’ and equations 10-11).** To this end, the KRT5/8 double-IF-stained sections used above for the assessment of cTEC network topology, were also utilized for mTEC morphometry development. To circumvent biases from potential inclusion of the epithelium-free regions corresponding to medullary PVS, we quantified the VI*_mTEC_* and II*_mTEC_* features, after excluding the PVS. Our analysis suggested that mTEC did not significantly decline with aging **(Figs. 6E-F)**. At first glance, this observation implies a relative resilience of the mTEC compartment, especially when compared to the cTEC. This outcome aligns with recent literature suggesting that aging predominantly affects the cTEC population, resulting in numerical and functional impairments, in turn, compromising thymopoiesis. In contrast, the impact of aging on mTEC is less profound^55,62,98^.

**Figure 6.**
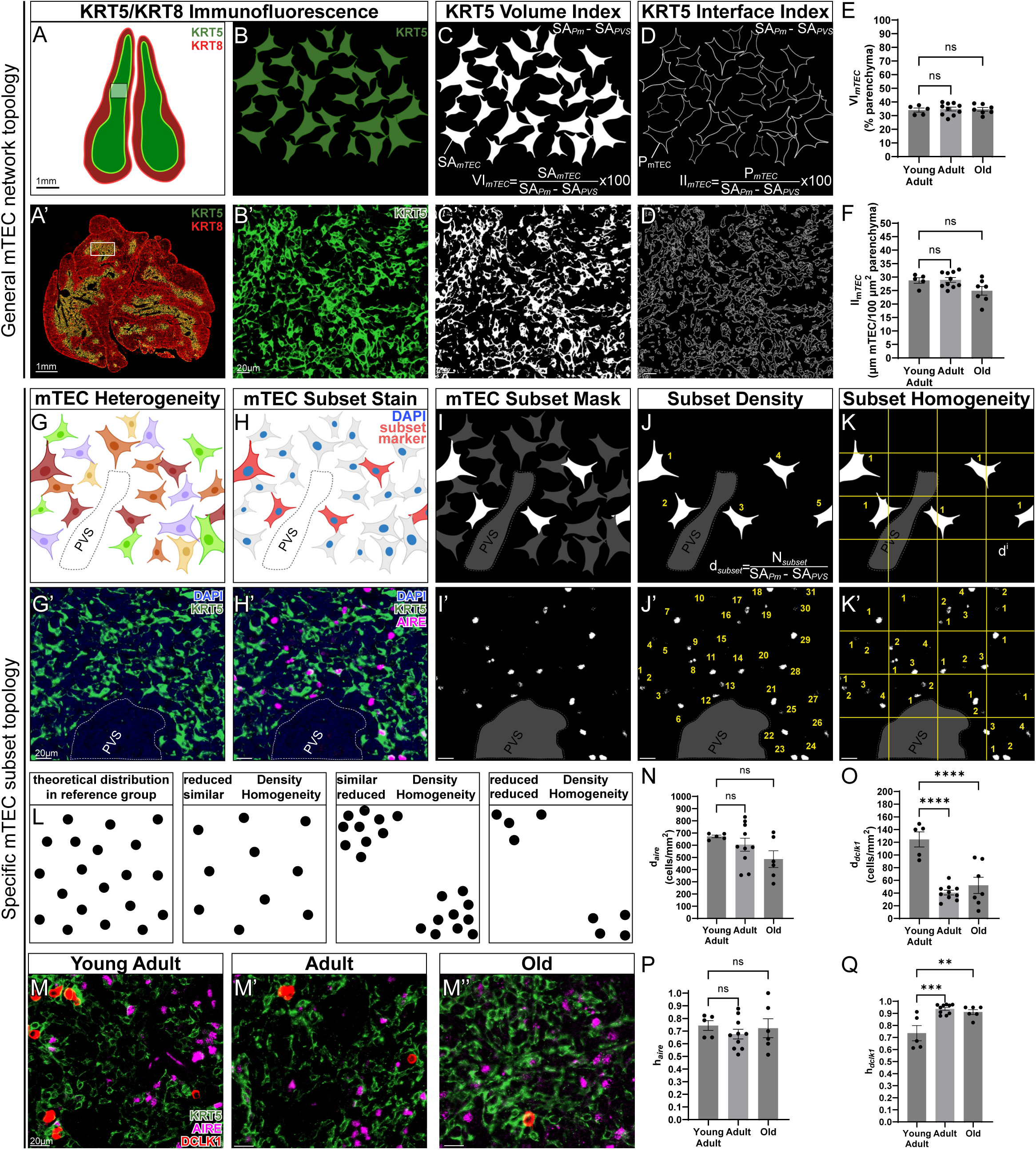
Topological assessment of the mTEC network. **(A-D)** Digital pathology pipeline for the topological assessment of mTEC network architecture. **(A-A’)** Whole thymi are stained for KRT5/8 immunofluorescence, and medullary regions are identified as KRT5^+^KRT8^+^. **(B-B’)** Thymic medulla field-of-view (FOV), depicting freeform arrangement of individual mTEC cells, although they are densely packed and close to one another. **(C-C’)** The binarization of the image shown in (B) via KRT5 signal intensity thresholding, creates a “mask” of the mTEC network. Accompanying formula for calculating the volume index, is shown to the bottom of the image (C). **(D-D’)** Development of outline from the binary mask shown in (C), revealing the interface area of the mTEC network. Accompanying formula for calculating the interface index, is shown to the bottom of the image (D). *Illustrations are shown in A-D, while actual staining is shown in the corresponding A’-D’ images.* **(E-F)** Morphometric assessment of mTEC network topology using mTEC volume index (VI*_mTEC_*) (E) and mTEC interface index (II*_mTEC_*) (F). *One-Way ANOVA; ns, not-significant.* **(G-K)** Digital pathology pipeline for the topological assessment of mTEC subset distribution. **(G-G’)** KRT5 staining reveals the mTEC network (G’), but not its heterogeneity, as implied by the multi-colored cells in (G). **(H-H’)** A special stain which is lineage-specific (H), such as AIRE (shown here in H’), or DCLK1 (not shown in this infographic), is used to label an mTEC subset of interest. **(I-I’)** A pixel classifier, or any other method, can be used to isolate the mTEC subset of interest (I), such as AIRE^+^ mTEC^hi^ (I’) from the rest of mTEC populations. **(J-J’)** Quantification of density of mTEC subset of interest, after quantification of individual cells within the field-of-view (J), such as AIRE^+^ mTEC^hi^ cells, is then normalized to the epithelium-containing medullary parenchyma (i.e., after excluding the PVS area from the parenchymal area of the medulla). Accompanying formula for calculating the density of mTEC subset of interest, is shown to the bottom of the image (J). **(K-K’)** Quantification of homogeneity of distribution of an mTEC subset of interest, through segmentation of a field-of-view in sectors, and assessing the density of the mTEC subset of interest in each individual sector (K), such as that of AIRE^+^ mTEC^hi^ within epithelium-containing medullary zones (K’). *Illustrations are shown in G-K, while actual staining is shown in the corresponding G’-K’ images.* **(L)** Theoretical adaptations of the distribution of an mTEC subset of interest, along with the expected changes in mTEC subset morphometries, following these adaptations. Black dots represent cells belonging to the mTEC subset of interest. **(M-M’’)** Representative images of medullary regions, stained with KRT5, DCLK1, and AIRE, to uncover mTEC subsets of interest, i.e., the DCLK1^+^ mTEC^hi^ tuft cells, and AIRE^+^ mTEC^hi^ cells in Young-Adult (M), Adult (M’), and Old (M’’) mice. **(N-O)** Quantification of the density of AIRE^+^ (d*_aire_*; N) and DCLK1^+^ (d*_dclk1_*; O) cells in the epithelium-containing medullary parenchyma, across different age groups. **(P-Q)** quantification of the homogeneity of AIRE^+^ (h*_aire_*; P) and DCLK1^+^ (h*_dclk1_*; Q) cell distribution in the epithelium-containing medullary parenchyma, across different age groups. *One-Way ANOVA; *p*≤*0.05; **p*≤*0.01; ***p*≤*0.001; ****p*≤*0.0001; ns, not-significant. Abbreviations: d_aire_/d_dclk1_, density of AIRE^+^/DCLK1^+^ cells; h_aire_/h_dclk1_, homogeneity of AIRE^+^/DCLK1^+^ cells; SA_mTEC_, surface area of mTEC mask; P_mTEC_, perimeter of mTEC outline; SA_Pm_ surface area of medullary parenchyma; SA_PVS_, surface area of perivascular space; VI_mTEC,_ volume index of mTEC; II_mTEC,_ interface index of mTEC*.

However, prior reports have demonstrated multiple functional mTEC impairments with aging, either by measuring the expression of key genes of mTEC functionality and differentiation, or by assessing mTEC progenitor activity^55^. Given that certain medullary functions, such as negative selection, are uniquely dependent on highly-specialized mTEC subsets, such as AIRE^+^ mTEC^hi^ and the newly discovered thymic mimetic cells, we reasoned that our topological investigations using digital pathology should additionally focus on mTEC subsets, instead of only on the entire mTEC network. To this end, we developed two non-redundant morphometric features of mTEC subset distribution, the density and the homogeneity of distribution.

The methods rely on implementing lineage markers that will help discern mTEC subset(s) of interest from background mTEC populations **(Figs. 6G-I)**. Here, we demonstrate the methods for two distinct mTEC subsets, the AIRE^+^ mTEC^hi^ subset, a master-regulator of negative selection^99–101^, and the post-AIRE DCLK1^+^ tuft cells, which belong to the thymic mimetic cells and mediate neuroendocrine and immunological functions in the medulla^102–105^. To distinguish these two mTEC subsets from other mTEC, we introduced antibodies, specifically targeting AIRE and DCLK1 **(Figs. 6G’-I’)**, which exhibited patterns of expected expression in the nucleus and the cytoplasm, respectively. Individual mTEC subsets were identified automatically, using a pixel/particle classifier **(Figs. 6I-I’)**. Next, the medullary parenchyma was demarcated following exclusion of the PVS, thus limiting all ensuing analyses to the epithelium-containing areas.

The first morphometric feature, i.e., density of AIRE^+^ (d*_aire_*) and DCLK1^+^ (d*_dclk1_*) mTEC, is expressed as AIRE^+^ and DCLK1^+^ mTEC numbers per mm^2^ of epithelium-containing medullary parenchyma **(Figs 6J-J’ and equations 12-13)**. For the second morphometry, i.e., homogeneity of AIRE^+^ (h*_aire_*) and DCLK1^+^ (h*_dclk1_*), each IF image is first segmented into multiple, equal, rectangular sectors and then, auto-fluorescent erythrocytes and other imaging artifacts are manually removed, before subjecting the images to homogeneity assessments **(Figs. 6K-K’, S6A-C and S6A’-C’)**. In each individual, artifact-corrected sector, the densities of AIRE^+^ and DCLK1^+^ mTEC are first calculated **(equations 14-15)**, and the homogeneity among sectors is then assessed **(equations 16-19)**. If the mTEC subset of interest was distributed in a homogenous arrangement within the medullary parenchyma, each sector would exhibit similar density and as such the homogeneity index for the respective mTEC subset would be high **(Fig. 6L)**. If, however, the mTEC subset of interest was heterogeneously arranged, i.e., it was arranged in small cellular clusters within the medullary parenchyma, then such cell clusters would stochastically fall within only a small portion of these sectors, thus leaving other sectors devoid of cells. The latter arrangement would result in dissimilar density profiles among the different sectors, and as such, a lower homogeneity index for the respective mTEC subset **(Fig. 6L)**. Based on this principle, the h*_aire_* and h*_dclk1_* calculate the fraction of the medullary sectors in each thymus, whose individual sector d*_aire_* and d*_dclk1_* fall within a single referent standard deviation from their mean density. Based on these assumptions **(equations 16-19)**, the h index gets values between “0” and “1”, with a “1” implying a perfectly homogeneous mTEC cell arrangement where all sectors present with similar cell density, while lower h values express varying degrees of clustered arrangement.

We assessed the morphometric features of mTEC subset distribution in our mouse model of age-dependent thymic involution. It has been previously demonstrated that the absolute number of AIRE^+^ mTEC^hi^ declines significantly during aging^106^. Using the d*_aire_* index, however, we found that their density remains unaltered during aging **(Fig. 6M-N)**. This observation implies that the numeric decline of AIRE^+^ mTEC^hi^ cells follows proportionally the reduction of the medullary volume, thus keeping a consistent density throughout aging. Moreover, there are no changes in the h*_aire_* index **(Fig. 6P)**, suggesting that aging does not impact their spatial arrangement either. Together, these observations indicate that despite the documented mTEC numerical decline^106^, the interactions between SP4/8 thymocytes and AIRE^+^ mTEC^hi^ cells are not essentially compromised from age-dependent intrathymic remodeling.

Finally, we performed similar investigations on DCLK1^+^ tuft cells, a thymic mimetic cell subset that has undergone further terminal differentiation from the AIRE^+^ mTEC^hi^ cells, thus overall belonging to the post-AIRE mTEC subsets^4,103,107^. Thymic tuft cells, similar to other mimetic cells in the thymus, present developing SP thymocytes with a unique subset of lineage-restricting tissue-restricted antigens, while AIRE^+^ exert broader functions, indicating complementarity of the two populations^15,102,103,107^. Interestingly, we uncovered a significant decline of thymic tuft cells with aging, as assessed through d*_dclk1_* measurement **(Figs. 6M&O)**. This decline began early at the Adult group, but did not reveal any signs of further decline in the Old mouse group **(Figs. 6M&O)**. Accompanying the density decrease, we strikingly found an increase in h*_dclk1_*, thus suggesting their sparser arrangement in the aged thymus **(Fig. 6Q)**. Being fewer in number than AIRE^+^ mTEC^hi^, thymic tuft cells attempt to achieve more strategic positioning via spatial remodeling during aging, to increase the probability of interacting with SP developing thymocytes. Together, these observations indicate that combining the density and homogeneity may provide detailed resolution of dynamic shifts of mTEC subsets in thymus disease.

## DISCUSSION

In this study, we developed a comprehensive digital pathology pipeline to analyze thymus microanatomy, combining multiple orthogonal approaches. Traditionally, research in the field has used population-based methods, such as flow cytometry, to study intrathymic dynamic changes, thus focusing on numerical ratios among cell subsets of interest^108^. However, these methods lack contextual information, such as the spatial distribution of stromal cells and thymocytes, within these complex thymic environments. Despite significant advancements in imaging methods, the microanatomy of the thymus remains underexplored, compared to other, particularly lymphoid, organs. Our pipeline circumvents this knowledge gap, offering insights into various pathologies, including acute or chronic thymic involution, and specific diseases affecting the thymus. Overall, our pipeline serves as a versatile tool for immunology labs specializing in thymus biology, homeostasis, and regeneration.

From development until preadolescence, the thymus generates a critical lifelong T cell repertoire to support systemic immunity, immune surveillance, and central tolerance^109–111^. However, it undergoes an age-dependent decline shortly after reaching the peak of functional maturity in adolescence. In late adulthood, systemic immunity and peripheral immunosurveillance becomes less dependent on the thymus, and more dependent on expansion of peripheral memory T cells^112–114^. Nevertheless, a recent epidemiologic study has challenged this premise^115^, suggesting that the thymus significantly contributes to systemic immunity even at older ages. Age-dependent thymic decline correlates with higher cancer incidence^5,116–118^, suggesting that targeting thymic involution could help prevent cancer and promote longevity^119^. Among thymic disorders, age-dependent involution is the most well-characterized thymic pathology^120^, and as such, served as the primary focus for demonstrating our pipeline’s utility. Additionally, the thymus is vulnerable to acute involution^9,121,122^, from factors like chemo/radiotherapy^123^, infections^121^, sexual steroids^124,125^, pregnancy^126,127^, and obesity^32,128^. Therefore, our pipeline may be utilized to offer insights in studies aimed at dissecting the molecular mechanisms and reversing thymic involution.

In addition, our digital pathology pipeline is a vital tool for studying thymic structure in various immunodeficiency and autoimmune disorders. For instance, MHC-II deficiency (MHCII-D), also known as Bare Lymphocyte Syndrome (BLS), a combined immunodeficiency syndrome due to mutations in genes regulating expression of MHC molecules^129^, is marked by increased corticomedullary ratio due to hypoplastic medulla, and a decline of AIRE^+^ mTEC^hi^ cells^130^. Similarly, AIRE deficiency leading to the autoimmune-polyendocrinopathy-candidiasis (APECED) syndrome, is characterized by loss of self-tolerance affecting multiple organs^100^. The syndrome is associated with medullary atrophy and diminished AIRE^+^ mTEC^hi^ cells^131^. Interestingly, it has been surmised that underlying the widespread loss of self-tolerance in peripheral tissues may be the dysregulation of terminally-differentiated post-AIRE subsets^132^. Of note, thymic tuft and myoid cells, two well-characterized thymic “mimetic” cell subsets, present unique chemosensory cell- and muscle cell-specific Tissue-Restricted Antigens (TRAs), respectively, to developing SP4/8 thymocytes, hinting on the APECED symptoms occurring in various endocrine organs and skeletal muscle^102,103,105,107,133,134^. A deeper understanding of the developmental pathways and spatial distribution of specific mTEC subsets, as demonstrated in our morphometric analyses, could thus offer insights on the onset and progression of primary immunodeficiencies, given that relevant mouse models are currently available^135–137^. In summary, our pipeline may be instrumental in exploring cTEC/mTEC adaptations in preclinical mouse models for rare immunological disorders.

A key feature of our morphometric approaches is their “orthogonality” as they offer alternate strategies for evaluating morphological characteristics of thymic environments and their cells. The most conspicuous example of orthogonality is the measurement of corticomedullary ratios, using either a “whole-thymus” or a “thymic-lobule” approaches. Interestingly, different imaging modalities within the same approach, i.e., IF or HE for the “whole-thymus” approach, revealed high correlation, suggesting they provide overlapping information. However, the same imaging modality, i.e., HE across different approaches: “whole-thymus” versus “thymic-lobule”, have been able to both capture the decreased corticomedullary ratio with aging, albeit via offering non-overlapping information, such as the status of thymic lobulation in the aged thymus. Although pathologists evaluate whole thymus slides using HE staining, according to the INHAND guidelines^1,26,28^, and the method is widely adopted in laboratories, whole-thymus approaches may suffer from significant limitations, such as sample orientation in the paraffin block, depth of sectioning, and tissue preservation, raising issues of reproducibility. Orthogonal approaches in morphometric measurements will thus provide high credibility of the reported data.

In imaging, “skeletonization” transforms binary objects to one-pixel-thick representations, facilitating topological analysis and reducing dimensionality. This technique, applicable to both 2D and 3D models of any branched network^138,139^, is notably used in studying vasculature development and angiogenesis^140^. Here, we adapted an ImageJ plugin^38^, to assess topological descriptors, such as number of branches/junctions, leveraging the meshwork architecture of the cTEC network, which resembles angiogenic vasculature^4,141^. To our knowledge, this is the first instance of using “skeletonization” to study the topology of the cTEC network in the murine thymus. Previous research documented the decline of the thymic cortex versus the medulla with aging, leading to decreased cTEC number and thymopoietic potential^28,111^ ^31^. Our findings in Old mice confirmed this decline via an “artificial” increase in cTEC network density, due to fewer thymocytes percolating through the network^31,111^. Interestingly, we additionally observed that the cTEC network undergoes hypertrophic growth in response to this decline, adjusting to the increased functional demands of thymopoiesis. The unique set of morphometric features we developed extracts key details of cTEC meshwork topology, revealing that cTEC adaptations relate to increased branching capacity without alterations in cell body thickness or tortuosity. Together, our pipeline not only corroborates earlier findings on age-dependent thymic changes, but also provides detailed insights into the nature of these changes. For instance, it unravels new possibilities for investigating cytoskeletal dynamics, as a putative strategy to tackle cortical adaptations in the aged thymus towards achieving better regenerative potential.

Multinucleation in TEC in both mice and humans has been sporadically reported. An older study using the C3H mouse strain found multinucleated cTEC and mTEC, with strong prevalence in the latter, with incidence increasing with age, and linked to thymic involution^142^. Similarly, binucleated and multinucleated cTEC were demonstrated in an acute thymic involution model, with mouse exposure to high altitudes, and confirmed in human patients^74^. These studies suggest that cTEC become multinucleated under certain (patho)physiological conditions, although the molecular prerequisites remain undetermined. In general, multinucleation arises either from cell fusion resulting in “syncytia”, or from nuclear division without cytokinesis resulting in “coenocytes”^143^. It has been speculated that multinucleation outside normal tissue development may enhance functional capacity, and facilitate rapid responses to high metabolic and regeneration demands^144–146^. Notable examples of this process are monocytes fusing into osteoclasts to facilitate bone remodeling^75^, and fused hepatocytes that promote liver regeneration^147^. Similarly, in the thymus, multinucleation might reflect a response to increased thymopoietic demands, which promote cTEC hypertrophy and syncytial configuration during involution. Electron microscopy has accurately displayed the potential of syncytial adaptation in thymic involution^74^. However, these studies are unable to probe the molecular basis of this adaptation, highlighting the need for future research to understand the functional decline of the aging thymus.

Thymocyte development relies on the presentation of MHC-self-peptide complexes by the cTEC/mTEC^148^. The wide repertoire of cortical self-peptides is orchestrated by unique proteolytic systems in cTEC, such as thymoproteosome β5t^77^, cathepsin-L^149^, thymus-specific protease PRSS16^150^, and the macroautophagy pathway^151^. Macroautophagy, a process capturing part of the cTEC/mTEC cytoplasm including organelles and vesicles for digestion in autophagolysosomes, is crucial for generating self-peptides, especially for MHC-II presentation^78,80,152^. Prior electron microscopy studies have identified autophagolysosomes in TECs, but the complexity of TEC heterogeneity and the specific identification of these vacuoles in different TEC subsets was not appreciated at the time^69,153^. Although we did not specifically explore the functions of this proteolytic system, we found that autophagolysosomes are strategically positioned both near the perinuclear cytoplasm and along cytoplasmic extensions surrounding thymocytes, possibly optimizing the sites for self-peptide loading. Age-dependent alterations in the number and content of these autophagolysosomes were evident, and possibly useful for extrapolating functional correlates of the cTEC network. Although not developed fully in our study, incorporating ultrastructural morphometry into our digital pathology pipeline has greatly enhanced our ability to identify the structural and functional deficits of the cTEC network with aging.

Upon successful negative selection, mature SP4/8 thymocytes leave the thymus trough a microanatomical compartment called the “perivascular space” (PVS)^17,39,85,86,88,94,96,154^. The PVS, which encircles medullary blood vessels, provides a distinct egress path, consisting of two concentric mesenchymal sheaths, an outer boundary made of adventitial fibroblasts and an inner boundary corresponding to the pericyte coating of the blood vessel^3^. To exit the thymus, SP4/8 thymocytes first migrate into the PVS passing through the first boundary, and then into the blood passing through the second boundary, following chemokine gradients, such as sphingosine-1 phosphate (S1P)^21,91,155–159^. The size and distribution of the PVS is therefore a direct functional correlate of thymic output^17^. Although it is barely visible in infants, the PVS significantly expands with age in humans, indicating egress defects in older individuals^94^. In our study, the age-related PVS expansion was observed using our novel morphometric analyses, demonstrating again, the orthogonality across different methods. Interestingly, the PVS serves as a niche for B cells and plasma cells, contributing to immune defense in healthy individuals^87^, but also to autoimmune responses under pathological conditions (e.g., Myasthenia gravis)^160^. Many of these conditions may critically affect PVS microanatomy without necessarily altering the overall corticomedullary architecture^132^, and as such, our morphometric approaches are suitable for providing insights on thymocyte emigration in thymic involution and disease.

Assessing both the density and homogeneity of various mTEC subsets within the medullary parenchyma is crucial for understanding the biological processes underlying intrathymic remodeling across different thymic pathologies, such as in the case of age-dependent decline. By analyzing these two metrics concurrently, we gained valuable insights into the intricate balance of mTEC subsets maintained within the aged thymus. Our findings revealed that while the density and homogeneity of AIRE^+^ mTEC^hi^ cells are relatively preserved, indicating a stable potential for interactions between AIRE^+^ mTEC^hi^ and SP4/8 thymocytes–and thus ensuring exposure to a broad spectrum of tissue-restricted antigens, there was notable decline in the density of DCLK1^+^ tuft cells, a post-AIRE thymic mimetic cell derivative. Interestingly, despite their reduced density, tuft cells demonstrate a more homogenous distribution within the aged thymus, possibly reflecting their strategic repositioning to optimize their interactions with SP4/8 thymocytes. This observation suggests a deliberate biological adaptation, favoring the preservation of AIRE^+^ mTEC^hi^ cells, which present a wider array of TRAs compared to that of tuft cells^15,102,103,107^. This strategic preservation of AIRE^+^ over DCLK1^+^ mTEC cells may be an evolutionary compromise to minimize the impact on central tolerance mechanisms and prevent the escape of self-reactive T cells^106^. Our study underscores the importance of considering both cell density and distribution patterns in understanding adaptive responses of the thymus to aging or other pathologies, and its implications for maintaining immune competence and preventing autoimmunity.

In conclusion, our comprehensive digital pathology pipeline not only offers a holistic view of the thymic environments with a broad spectrum of morphometric analyses, but also serves as a practical toolset for other laboratories interested in thymus pathology. This integrated approach enables the detailed exploration of thymic architecture and its adaptations to aging and disease, providing a platform for the study of cellular distributions, density, and structural dynamics. By offering a versatile and accessible resource, our work facilitates unified investigations into thymic biology, paving the way for novel therapeutic strategies aimed at enhancing thymic regeneration and bolstering immune resilience across the lifespan.

## Supporting information

Lagou et al. Supplementary Material

## ACKNOWLEDGEMENTS

We thank Dr. Ana-Maria Cuervo for providing the 2-year-old C57BL/6J mice for our studies. We also thank the Histopathology Core for processing and paraffin-embedding of formalin-fixed thymic and splenic samples.

Several investigators would like to acknowledge the following for funding support: the new investigator start-up funds, provided to G.S. Karagiannis from the Montefiore-Einstein Comprehensive Cancer Center (P30 CA013330); the Careers in Immunology Fellowship (2022), awarded to M.K. Lagou and G.S. Karagiannis from the American Association of Immunologists; the American Society of Hematology Junior Faculty Scholar Award (2023), provided to M. Maryanovich; the NIH T32 HL144456 (NHLBI), the NIH F32 HL158084 (NHLBI), the Gottesman Stem Cell Institute Paul S. Frenette Scholars Award, and the Cancer Research Institute Irvington Postdoctoral Fellowship (CRI4994), provided to R.S. Carpenter.

We would also like to acknowledge the use of the Zeiss Supra 40 Field Emission Scanning Electron Microscope (1S10RR025554-01A1) and the 3DHISTECH P250 High-Capacity Slide Scanner (1S10OD026852-01A1) in the Analytical Imaging Facility (AIF), both acquired through Shared Instrumentation Grants. The AIF was also partially funded by the NCI Cancer Center Support grant (P30CA013330).

Author contributions: M.K. Lagou and G.S. Karagiannis designed the study; M.K. Lagou, D.G. Argyris, S. Vodopyanov, L. Gunther-Cummins and C. Entenberg performed experiments; L. Gunther-Cummins, H. Guzik, X. Nishku, J. Churaman performed sample preparation for all the imaging studies; M.K. Lagou, D.G. Argyris, S. Vodopyanov, L. Gunther-Cummins, S. DesMarais, C. Entenberg, R.S. Carpenter, V. DesMarais, and F.P. Macaluso performed image analysis; A. Hardas, and T. Poutahidis performed reviews of all thymic and splenic slides and diagnostic pathology reviews on the spontaneous lesions of aged mice; M.K. Lagou, D.G. Argyris, C. Panorias, M. Maryanovich, and G.S. Karagiannis performed data analyses and statistical analyses. M.K. Lagou, D.G. Argyris, S. Vodopyanov, and G.S. Karagiannis wrote the final report. All authors contributed to the editing of the final report. All authors agree to the content of the submitted manuscript.

All figures were created with BioRender.com and Abode InDesign and Illustrator.

