## Supplementary material for "Morphometric Analysis of the Thymic Epithelial Cell (TEC) Network Using Integrated and Orthogonal Digital Pathology Approaches": Lagou et al. Supplementary Material

### Supplementary Figure 1

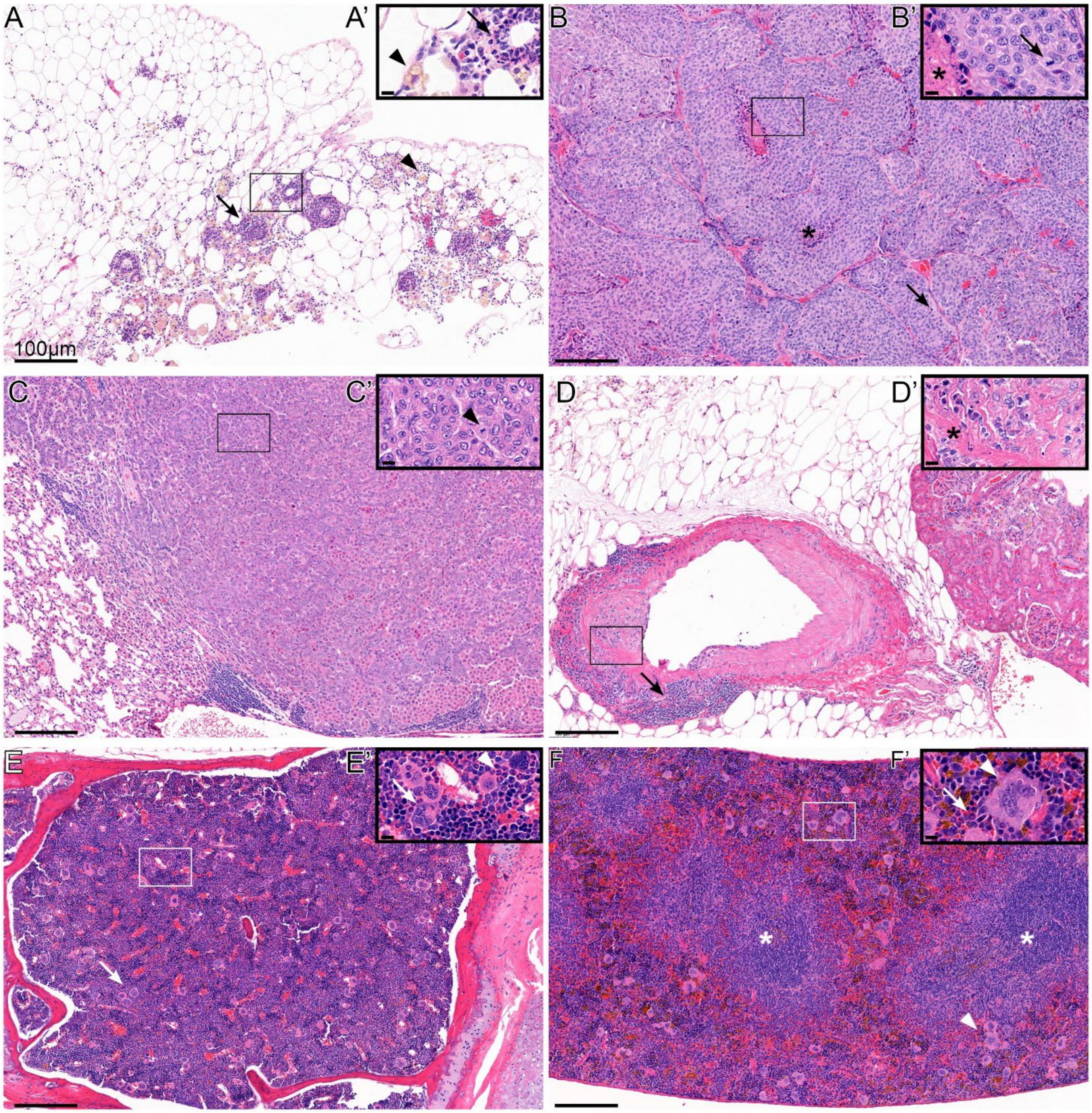


**Supplementary Figure 1. (A-H)** Incidental histopathological lesions in Old mice. **(A-A') Omentum:** multifocal mild-to-moderate neutrophilic, histiocytic, lymphoplasmacytic, necrotising steatitis (arrows) with hemosiderin laden macrophages (arrowheads). **(B-B') Adrenal Gland:** malignant pheochromocytoma. Neoplastic cells are arranged in variably sized, irregular lobules separated by thin collagenous septa. Lobules often show central necrosis (asterisks). Moderate-to-high number of mitoses (arrows). **(C-C') Lung:** solid, bronchiolo-alveolar carcinoma. Atypical round-to-cuboidal cells with eosinophilic cytoplasm and large, round euchromatic nuclei are closely packed with minimal arrangement of stromal elements. Arrowhead depicts mitotic figure. **(D-D') Perirenal Fat Vessel:** chronic vasculitis. The vessel wall is expanded by a mixed infiltrate of lymphocytes (arrows), and fewer neutrophils and cellular debris. Asterisk depicts vessel wall necrosis. **(E-E') Bone Marrow, Sternum:** The cellularity of granulocytic lineages is increased, with variable myeloid cells filling the medullary space, and increasing the myeloid-to-erythroid ratio. Prominent megakaryocytes are also present (arrows). Arrowhead depicts granulocytic precursor cells. **(F-F') Spleen:** Throughout the splenic parenchyma (red pulp) are numerous megakaryocytes (arrowheads) and moderate numbers of erythroid and myeloid precursors (arrow) with occasional mitotic activity (extramedullary hematopoiesis). Splenic architecture is maintained with evident lymphoid follicles (asterisks). There are multifocal increased numbers of hemosiderin$^{+}$ macrophages (brown pigment). *Scale Bars: A-F, 100μm; A'-F', 20μm*

### Supplementary Figure 2

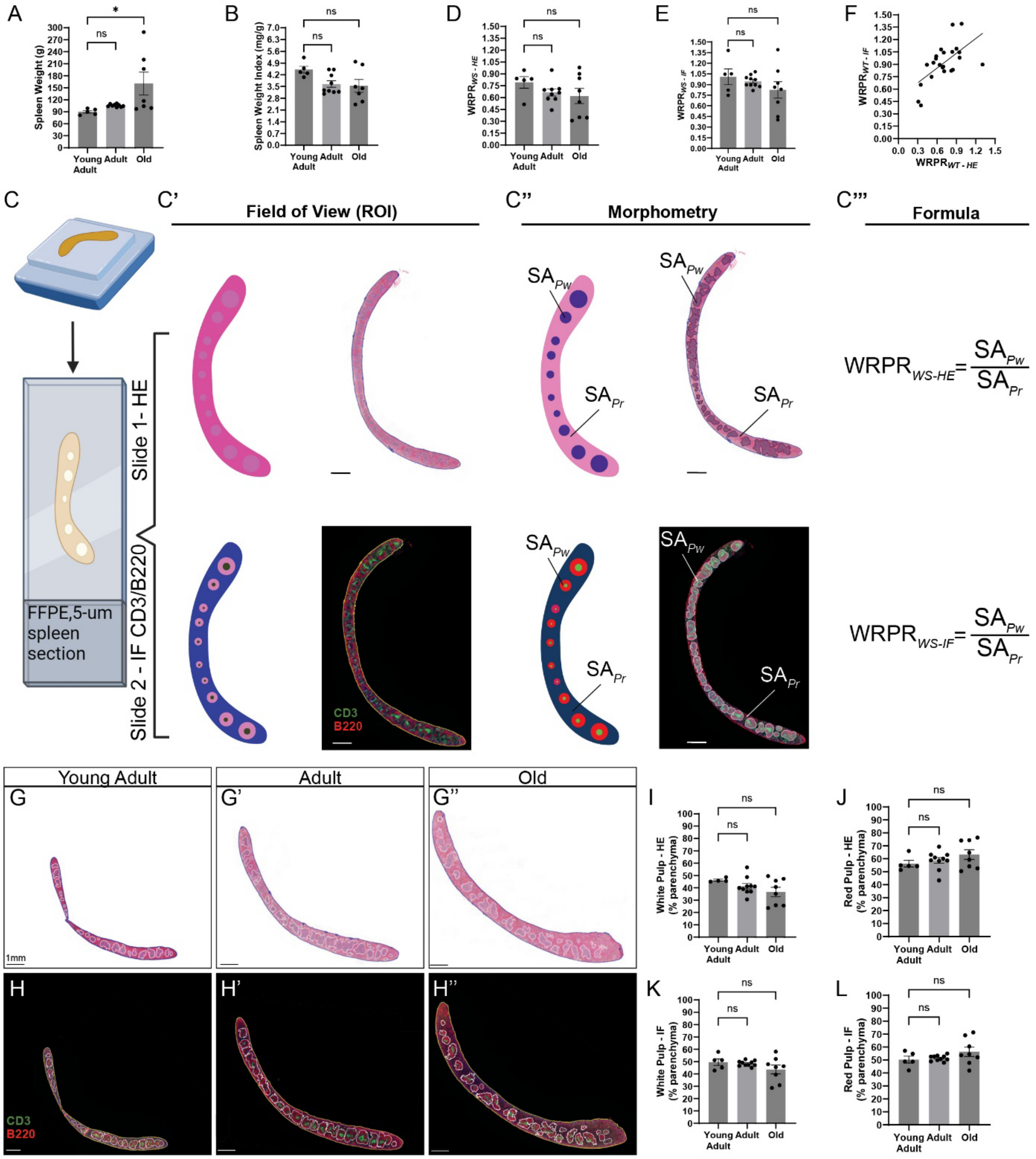


**Supplementary Figure 2. White-to-Red Pulp Ratio (WRPR) Assessment. (A)** Absolute Spleen Weight (g) in the mouse cohort. **(B)** Spleen Weight Index (SWI, mg of spleen per gr of body weight). **(C-C''')** Pipeline for assessing WRPR in whole-spleen slides ($WRPR_{WS}$), using HE (slide 1) and IF (slide 2) methods. 5µm thick sections from FFPE blocks are HE and IF stained (C). First, the entire splenic regions are annotated on the HE and IF slides (blue and yellow demarcation, respectively) (C'), followed by the white pulp (purple and white annotations, respectively) (C''). Respective WRPR Formulas are shown in (C'''). **(D-E)** WRPR quantification, using the HE (D) or the IF (E) method, in mice across different age groups. **(F)** Correlation plot with simple linear regression line between the HE and IF methods of calculating WRPR. **(G-H)** Whole-slide images of spleens using the HE (G-G''), or the IF (H-H'') method, representative of Young-Adult (G&H), Adult (G'&H') and Old (G''&H'') mice. **(I-J)** Surface Area percentage (%) of the white (I) and red (J) pulp in the HE sections. **(K-L)** Surface Area percentage of the white (K) and red (L) pulp in the IF sections. *Abbreviations: HE, Hematoxylin-Eosin; IF, Immunofluorescence; FFPE, Formalin-Fixed Paraffin-Embedded; SWI, Splenic Weight Index; $SA_{Pw}$, Surface Area of White Pulp; $SA_{Pr}$, Surface Area of Red Pulp; $WRPR_{WS}$, White-to-Red Pulp Ratio of the whole spleen. One-Way ANOVA, (*) p≤0.05, ns: no significance.*

**Supplementary Figure 3**

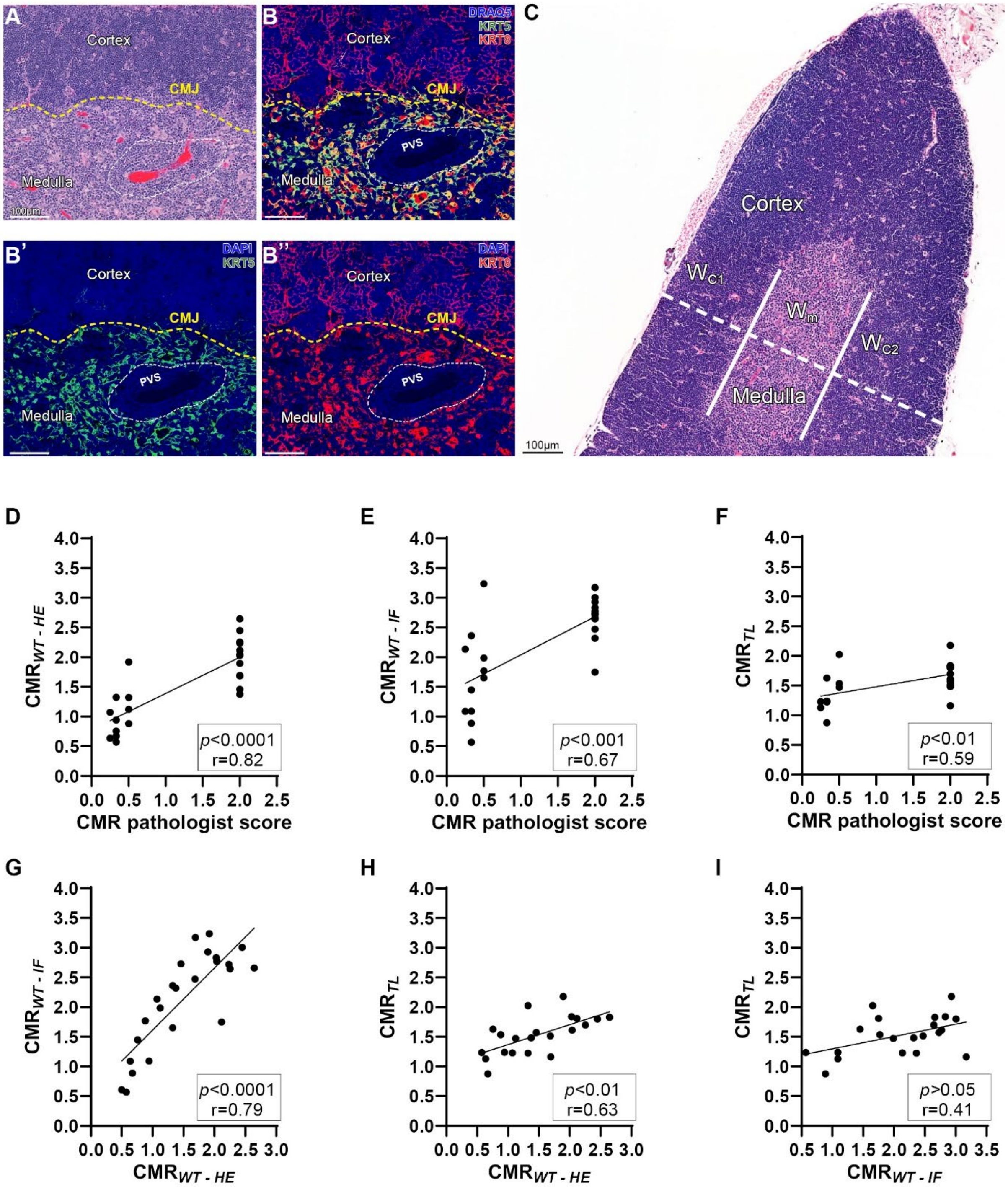


**Supplementary Figure 3. Assessment and Validation of Corticomedullary Ratio (CMR) Morphometry.** **(A-B)** Representative images of the CorticoMedullary Junction (CMJ) from a Young-Adult mouse. The cortex is distinguished from the medulla by the higher density of hematoxylin in thymocytes (A; HE section), or by $KRT8^+KRT5^-$ keratin profile (B-B''; IF section); in contrast, the Medulla is characterized by $KRT8^+KRT5^+$ area (B-B''). **(C)** Representative thymic lobule using an HE section that fulfills the criteria for quantification of the $CMR_{TL}$ morphometry, i.e., the vector segmenting the lobule into cortical and medullary segments (white-dotted line) is perpendicular to the two CMJ borders (white lines) **(D-I)** Correlation plots with simple linear regression line between the four different methods, including the pathologist-based method, of calculating CMR, as shown in the correlation matrix in the Figure 2I of the manuscript. *Abbreviations: HE, Hematoxylin-Eosin; IF, Immuno-fluorescence; KRT5, Keratin 5; KRT8, Keratin 8; CMR, CorticoMedullary Ratio; PVS, PeriVascular Space;* $W_{c1}/W_{c2}$*, Width of Cortex Segment 1 and 2;* $W_M$*, Width of Medulla Segment. Scale Bars, 100µm. Spearman's correlation coefficient of determination: (*), p≤0.05; (**), p≤0.01; (***), p≤0.001; (****), p≤0.0001.*

### Supplementary Figure 4

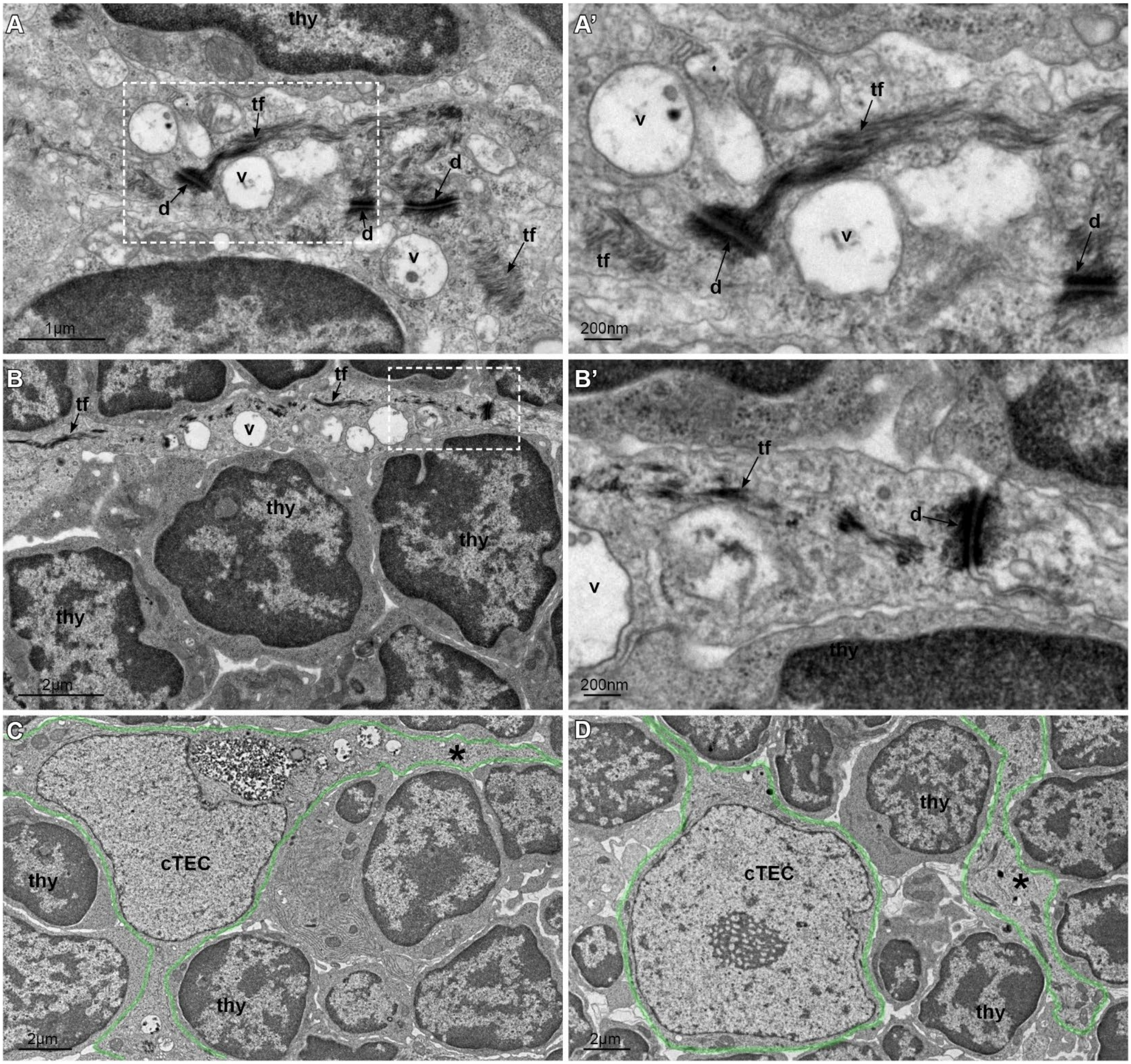


**Supplementary Figure 4. Ultrastructural Topography of the Cortical Parenchyma and the cTEC Meshwork. (A-B)** Identification of cTEC cytoplasmic segments travelling among thymocytes (thy). All segments lack an immediate connection to a discernible cTEC within the same viewing plane, as characterized by its euchromatic nucleus. This identification utilizes three key ultrastructural features, i.e., desmosomes (d), cytoplasmic vacuoles (v), and tonofilaments (tf), which are all exclusive of thymic epithelial cells within the thymic parenchyma. Panels (A') and (B') are magnified areas highlighted by the white-dotted lines in panels (A) and (B), respectively. **(C-D)** Representative images, where cTEC cytoplasmic segments appear to be proximal to a characteristic cTEC nucleus in the same viewing plane. The identification is easier, but it still employs the same ultrastructural features for identification. In panel (C), the segment (asterisk) appears to extend as a "continuum" with the perinuclear cTEC cytoplasm, whereas in panel (D), the segment (asterisk) shows no direct continuation with the perinuclear cTEC cytoplasm, thus raising the possibility that it may or may not be part of this particular cTEC. This figure suggests that identification of key cTEC features (i.e., d, v, and tf), is crucial for ultrastructural topography, and should not be considered redundant.

**Supplementary Figure 5**

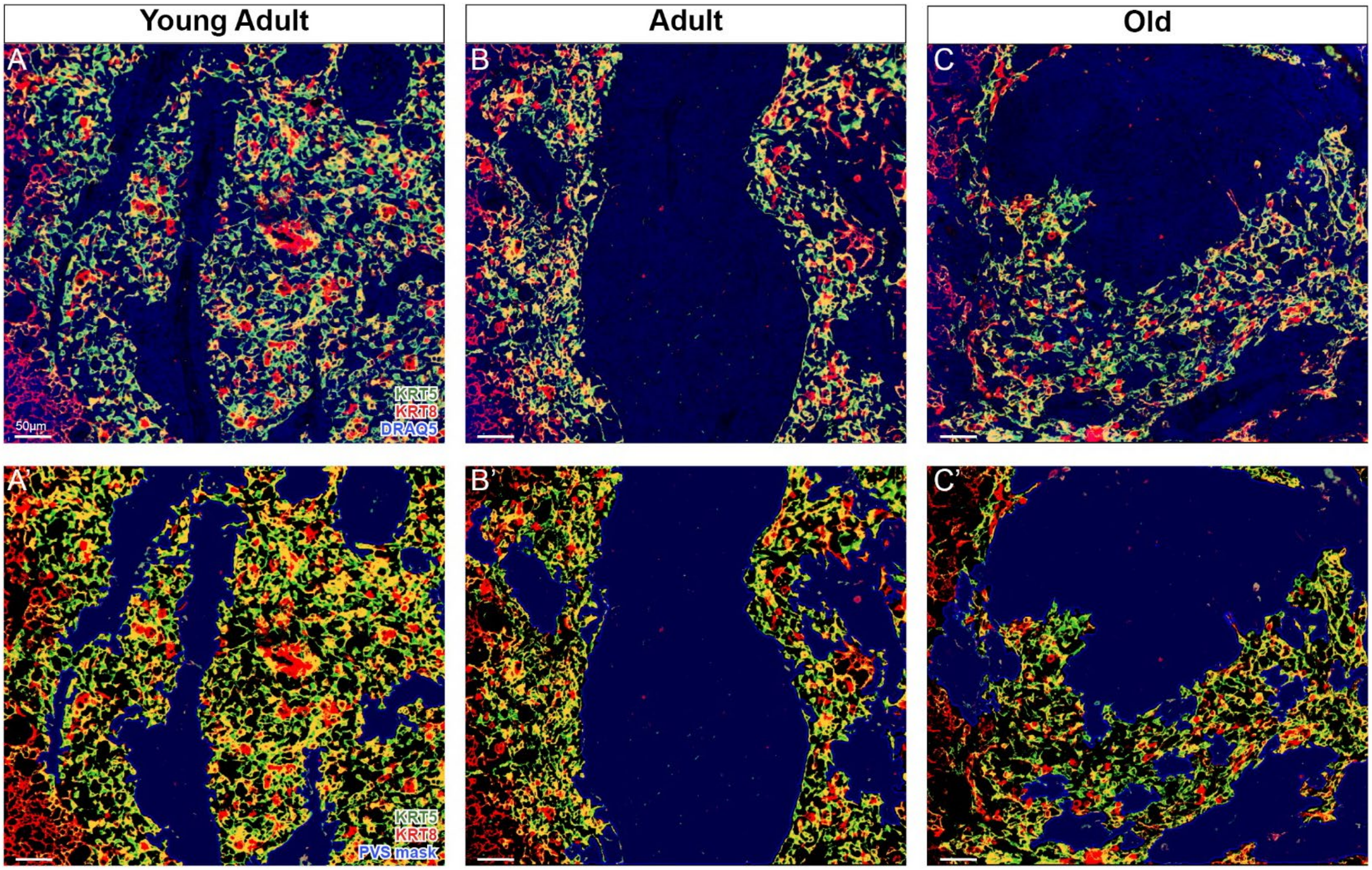


**Supplementary Figure 5. (A-C & A'-C') Perivascular Space in Mouse Thymus Across Age Groups.** Expanded views from magnified inserts depicted in Figures 4D-F of the manuscript. The figure includes images without the Perivascular Space (PVS) mask annotation (A-C), thus highlighting the presence of cellular components, specifically DRAQ5$^{+}$ nuclei, within the PVS regions. Additionally, images with the PVS mask annotation (A'-C') are provided, as illustrated in the original figures, for comparisons. The overall purpose of this figure is to affirm that the PVS regions quantified in our morphometric analysis are indeed populated with cells, and do not represent tissue processing artifacts or void spaces.

### Supplementary Figure 6

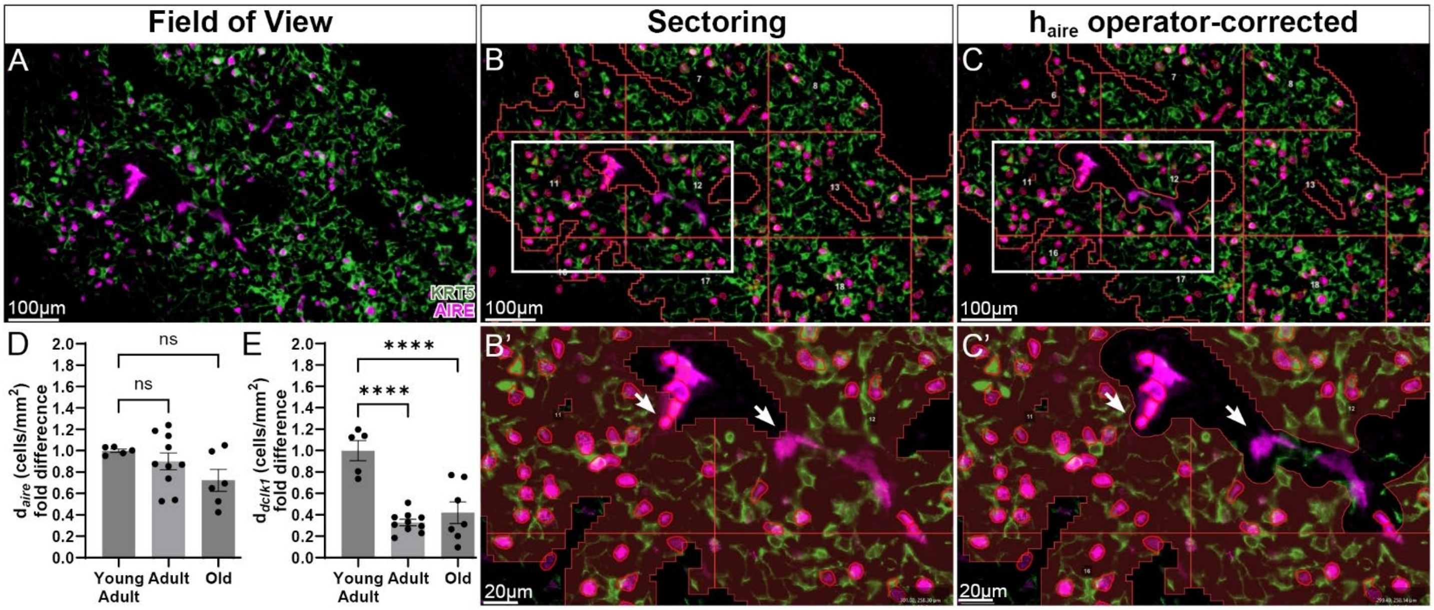


**Supplementary Figure 6. Topological Features of mTEC Subset Distribution in Medullary Parenchyma. (A-C)** Digital pathology pipeline for the development of two features of mTEC subset distribution, density (d), and homogeneity (h) of distribution. A medullary region is initially captured as field-of-view (A), demarcating the epithelium-containing area (ECA) through generic KRT5 staining to label all mTEC, as well as IF staining of a lineage marker of interest, such as AIRE, which demarcated the $mTEC^{hi}$ subset regulating negative selection. Automatic algorithms, as described in the manuscript's methods, are then applied to segment the field-of-view into multiple equal sectors (their borders shown as yellow lines), after excluding the epithelium-free perivascular space. This algorithmic implementation results in the development of multiple sectors with irregular shape and different surface area of $KRT5^+$ medullary parenchyma (B). A pixel classifier is then utilized to quantify the $AIRE^+$ cells in each sector independently (B). Panel B' represents a magnification of the white box shown in panel B, showing an intersection point among multiple sectors. As the last step of this process, artifacts such as autofluorescence etc., especially intravascular erythrocytes that provide $AIRE^+$ false-positive signal (arrows), are removed from the image, using manual drawing tools (C-C'). **(D-E)** Normalized density of $AIRE^+$ (D) and $DCLK1^+$ mTEC, across different age groups. Because normalization was done with the mean of the control group, i.e., Young-Adult, individual values in these scatterplots reflect to fold-change from the mean of the Young-Adult group. *Abbreviations: $d_{aire}$, density of the $AIRE^+$ mTEC subset; $d_{dclk1}$, density of the $DCLK1^+$ mTEC subset. Scale Bars, 100µm. One-way ANOVA; (*), p≤0.05; (**), p≤0.01; (***), p≤0.001; (****), p≤0.0001; ns, no significance.*
